# Flow-Dependent CFU Dynamics Reshape Polymicrobial Biofilm with a Pronounced Dominance Shift under Sub-Inhibitory Antibiotic Stress

**DOI:** 10.64898/2026.08.18.745464

**Authors:** Ichha Viren Shah, Riddhin Modi, Devarshi Gajjar

**Affiliations:** Department of Microbiology and Biotechnology Centre, Faculty of Science, The Maharaja Sayajirao University of Baroda, Vadodara 390002, Gujarat, India

**Keywords:** CAUTI, polymicrobial biofilm, sub-MIC antibiotics, microfluidics, species dominance, flow conditions, Ciprofloxacin, Gentamicin

## Abstract

Catheter-associated urinary tract infections (CAUTIs) are the most prevalent healthcare-associated infections globally, yet the ecological dynamics governing polymicrobial biofilm communities on indwelling catheters remain poorly understood under physiologically relevant conditions. Most prior work uses static *in vitro* models that fail to capture continuous urine flow and sub-inhibitory (sub-MIC) antibiotic gradients. We investigated how continuous flow and sub-MIC concentrations of ciprofloxacin and gentamicin reshape colony-forming unit (CFU) dynamics across attached biofilm and dispersed effluent fractions, and species dominance in mono- and polymicrobial biofilms of *Pseudomonas aeruginosa* (Pa), *Klebsiella pneumoniae* (Kp), and *Enterococcus faecium* (Ef) using silicone-coated latex catheter segments, volumetric infusion pumps, and ibidi µ-slide VI ^0.4^ microfluidic chambers. Under antibiotic-free conditions, Pa dominated both dual co-cultures (Pa+Kp, Pa+Ef) in static condition, but this dominance was not sustained under flow in the Pa+Ef pairing, where Ef rose to 62.5% relative abundance. Sub-MIC ciprofloxacin under flow promoted Kp dispersal (+15.87 log₂ fold change in dispersed-cell fraction(filter), cooperative Pa recovery via Ef co-occupancy, and pronounced Ef dominance in the triple-species community (64.71% relative abundance). Ef exhibited enhanced growth under sub-MIC gentamicin in static conditions that was abolished under flow. CLSM imaging revealed ciprofloxacin-induced Kp filamentation under flow, with Ef microcolonies localising at filament termini—a novel architectural interaction providing spatial scaffolding for the gram-positive partner. These findings establish that continuous flow and antibiotic class jointly determine polymicrobial dominance outcomes in ways invisible to static assays, underpinning Ef persistence in mature CAUTI biofilms and highlighting flow as a central ecological variable in infection pathogenesis.

**IMPORTANCE:** CAUTIs account for up to 40% of all hospital-acquired infections worldwide, yet treatment frequently fails because catheter biofilms are polymicrobial—a reality invisible to single-species static models. This study subjects three-species communities of *P. aeruginosa, K. pneumoniae,* and *E. faecium* to continuous fluid flow and subinhibitory antibiotic concentrations that mimic conditions in a real urinary catheter. The present study demonstrates that flow rewires the competitive hierarchy, elevating *E. faecium* to community dominance under conditions where static assays predict its exclusion. We further demonstrate that sub-MIC ciprofloxacin induces *K. pneumoniae* filamentation that generates novel attachment surfaces exploited by *E. faecium*—a form of antibiotic-driven architectural remodelling with direct implications for treatment failure. These findings demand a fundamental reappraisal of CAUTI biofilm research and antibiotic dosing strategies for polymicrobial infections in catheterised patients

## INTRODUCTION

Catheter-associated urinary tract infections (CAUTIs) are the most prevalent category of healthcare-associated infections (HAIs) globally, accounting for up to 40% of all nosocomial infections (1, 2). Approximately 70% of UTIs arising in hospitalised patients are directly attributable to indwelling urinary catheters, with device utilisation ratios of 0.5–0.86 per 1,000 catheter-days in resource-constrained settings and polymicrobial biofilm communities that form on the luminal and extraluminal surface of the indwelling catheters within 24-48 h of insertion (3, 4). Biofilm-resident organisms are recalcitrant to both antimicrobial therapy and host immune clearance, producing chronic or recurrent infection that prolongs hospitalization, drives antimicrobial resistance selection, and accounts for substantial morbidity and mortality in intensive care settings (3, 5).In Indian intensive care units, CAUTI rates are reported to significantly exceed international benchmarks, driven by multidrug-resistant (MDR) gram-negative organisms — principally MDR *Klebsiella pneumoniae* (32%) and *Acinetobacter baumannii*, exhibiting >70% resistance to carbapenems and cephalosporins (3, 6).

While operationally tractable, existing CAUTI biofilm research fundamentally relies on static *in vitro* models whose quiescent conditions eliminate continuous urine flow and physiological shear rates, imposing assumptions inconsistent with the catheterized urinary tract. *In vivo*, catheter surfaces are exposed to continuous low-velocity urine flow that generates shear forces shaping biofilm architecture, selecting for adherent phenotypes, modulating nutrient gradients (i.e. nutrient replenishment and metabolic waste removal that sustain deep-layer microbial activity *in vivo*), and determining which species can establish and maintain surface colonisation (7, 8). As a consequence, static assays systematically misattribute competitive dominance to strong surface-adhering species, particularly Gram-negative rods such as *P. aeruginosa*, and cannot capture the ecological dynamics such as dispersal, redistribution, competitive exclusion, protective coexistence; that determine the community composition of a mature catheter biofilm exposed to antibiotic treatment over clinically relevant time scales (8, 9).

A further dimension of clinical relevance is the exposure of biofilm communities to sub-inhibitory concentrations (sub-MIC) of antibiotics. Limited drug penetration through the extracellular polymeric substance matrix, sequestration by anionic polysaccharides, and pH-mediated inactivation ensure that bacteria in the interior of catheter biofilms rarely encounter drug concentrations approaching the MIC determined for planktonic cells (10, 11).

Sub-MIC antibiotic exposure is not merely a state of incomplete inhibition. It is an ecological stressor with metabolic impacts that can induce the over expression of biofilm genes, stimulation of EPS, enhance persister cell formation, SOS activation and resistance mutation selection (12–14). Critically, these effects have been studied almost exclusively in monoculture, leaving the consequences for polymicrobial community structure and dominance hierarchies largely unexplored.

There has been limited systematic investigation of flow’s ecological effects on polymicrobial biofilm composition. According to previous studies using flow-cell and microfluidic biofilm systems, continuous shear imposes strong selection on biofilm architecture. The physical selection promotes compact, surface adherent phenotypes in strongly attaching species and the dispersal of weakly adherent populations (15, 16). Microfluidic devices allow for precise control of fluid shear, nutrient concentration, and antibiotic gradients. This allows for experimental conditions that cannot be achieved with static systems (17).

However, flow-cell and microfluidic biofilm studies have predominantly focused on monoculture systems (18). Those that have used mixed-species communities have rarely incorporated the clinically relevant context of simultaneous sub inhibitory antibiotic stress. As a result, the fundamental question of whether flow conditions alter the identity of the dominant competitor within a polymicrobial catheter community particularly under simultaneous antibiotic challenge remains unanswered.

*Enterococcus faecium* presents a particularly compelling ecological paradox in catheter biofilm biology. It is consistently recovered from polymicrobial CAUTI biofilms and represents over 50% of *enterococcal* clinical isolates in many tertiary care hospitals (19, 20). Its hospital-adapted CC17 lineage, which encompasses the majority of vancomycin-resistant *E. faecium* (VREfm) strains, is remarkable for its persistence in long-term catheterized patients despite a characteristically weak capacity for abiotic surface adhesion in monoculture (20). Biofilm-forming assays consistently classify *E. faecium* as a weak producer compared to *P. aeruginosa* or *K. pneumoniae*, leading to the inference that it is a minor or incidental member of polymicrobial catheter communities. Yet clinical and metagenomic evidence contradicts this inference (21). *K. pneumoniae* and *P. aeruginosa* are both ESKAPE pathogens on the WHO Bacterial Priority Pathogens List, with carbapenem-resistant designated high priority (22). They are recovered at substantial frequency from catheterized patients, with *Pseudomonas* implicated in up to 16% of ICU-associated UTIs (23) and *Klebsiella* recovered across independent CAUTI cohorts spanning multiple countries and care settings (24, 25). Unlike *E. faecium*, *Klebsiella* is an established strong biofilm-former on catheter surfaces under both static and flow conditions individually (26). *Pseudomonas* shows comparably strong individual biofilm-forming capacity and has been characterized in dual-species systems although almost always under static conditions. There are flow-based studies involving dual-species of *Pseudomonas*, such as those with *Candida albicans* (21, 27) and *Proteus mirabilis* (28, 29) . However, polymicrobial context for both these species under flow and in presence of sub-inhibitory antibiotic exposure remains unexamined.

In this study, we used a catheter-segment flow model with volumetric infusion pumps and ibidi µ-slide VI ^0.4^ microfluidic chambers to systematically characterize how physiologically relevant continuous flow and sub-inhibitory ciprofloxacin or gentamicin exposure restructure polymicrobial communities comprising *P. aeruginosa*, *K. pneumoniae*, and *E. faecium* clinical CAUTI isolates stratified across distinct antimicrobial susceptibility distributions. We combined CFU enumeration on chromogenic selective agar with confocal laser scanning microscopy (CLSM) viability imaging to resolve both abundance and spatial architecture

## RESULTS

### Biofilm and Adhesion Assay

All clinical isolates of *P. aeruginosa* (n=5) were classified strong biofilm producers and most of *K. pneumoniae* (n=5) isolates were classified moderate biofilm producers, whereas all *E. faecium* isolates (n=5) were classified weak biofilm producers (Figure 1a). Mixed-species combinations (Pa+Kp, Pa+Ef, Kp+Ef, Pa+Kp+Ef) consistently formed strong biofilms, though absolute OD₅₉₀ values were reproducibly lower than *P. aeruginosa* monocultures (Figure 1b). Indicating that co-culture attenuates the dominant Pa biofilm phenotype rather than enhancing it.

**Figure 1.**
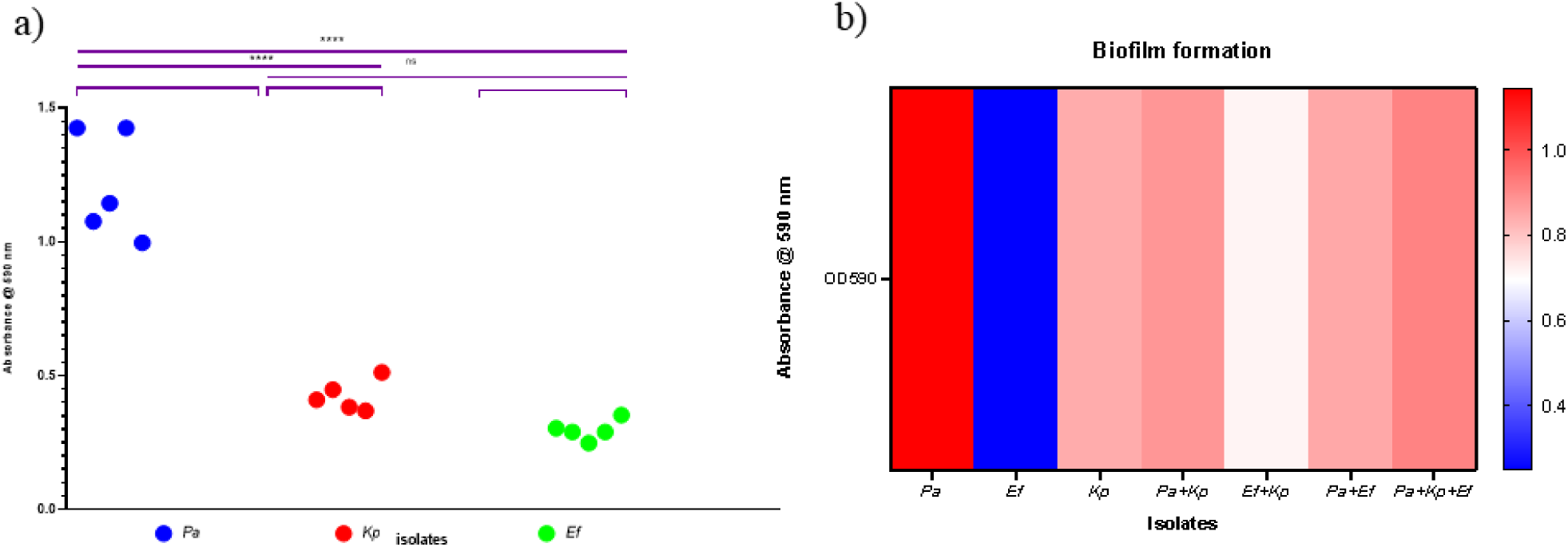
Quantification and categorisation of biofilms: a) Nested scatter plot showing crystal violet biofilm formation (OD590) for all clinical isolates of *Pseudomonas aeruginosa* (Pa; n = 5), *Klebsiella pneumoniae* (Kp; n = 5), and *Enterococcus faecium* (Ef; n = 5). All Pa isolates were strong biofilm producers; Kp isolates were moderate biofilm producers; all Ef isolates were weak biofilm producers. b) quantification of biofilm of representative isolates and its combination sets. Blue represents weak biofilm formers while red represent strong biofilm formers. Statistical analysis by one-way Anova and graph were made using GraphPad Prism 8.0 with p ≤ 0.01 (*), p ≤ 0.001 (**), and p ≤ 0.0001 (***).

In the adhesion assay, *P. aeruginosa* and *K. pneumoniae* exhibited a dense, uniform monolayer of adhered cells, whereas *E. faecium* displayed sparse, dispersed surface coverage (Figure 2-i A-C). Ciprofloxacin at sub-MIC exposure dramatically reduced adherent cell density across all monocultures (Figure 2-ii A-C), notably Kp displayed elongated rod morphology consistent with SOS-induced filamentation (Figure 2-ii B). Gentamicin produced a more moderate reduction in adhesion for Pa and Kp, while Ef adhesion remained comparable to untreated controls (Figure 2-iii A-C). In Under sub-MIC gentamicin, *E. faecium* became proportionally more prominent in Kp+Ef combinations (Figure 2-iii F) and Pa+Ef combinations (Figure 2-iii E) were preserved or increased relative to untreated controls, suggesting intrinsic gentamicin tolerance providing a relative competitive advantage and microcolonies were distinct in the latter. In polymicrobial conditions, Pa+Kp (Figure 2-i D) exhibited reduced adhesion compared to their respective monoculture, suggesting competitive interaction, while the triple species combination (Figure 2-i G) demonstrated robust surface colonization. In Pa+Ef co-culture (Figure 2-i E) without antibiotics, enhanced rod adhesion was observed relative to Pa monoculture, with only limited cocci visible, suggesting Pa forms an initial scaffolding that stabilises Ef surface retention despite the latter’s inherently weak adhesion capacity.

**Figure 2.**
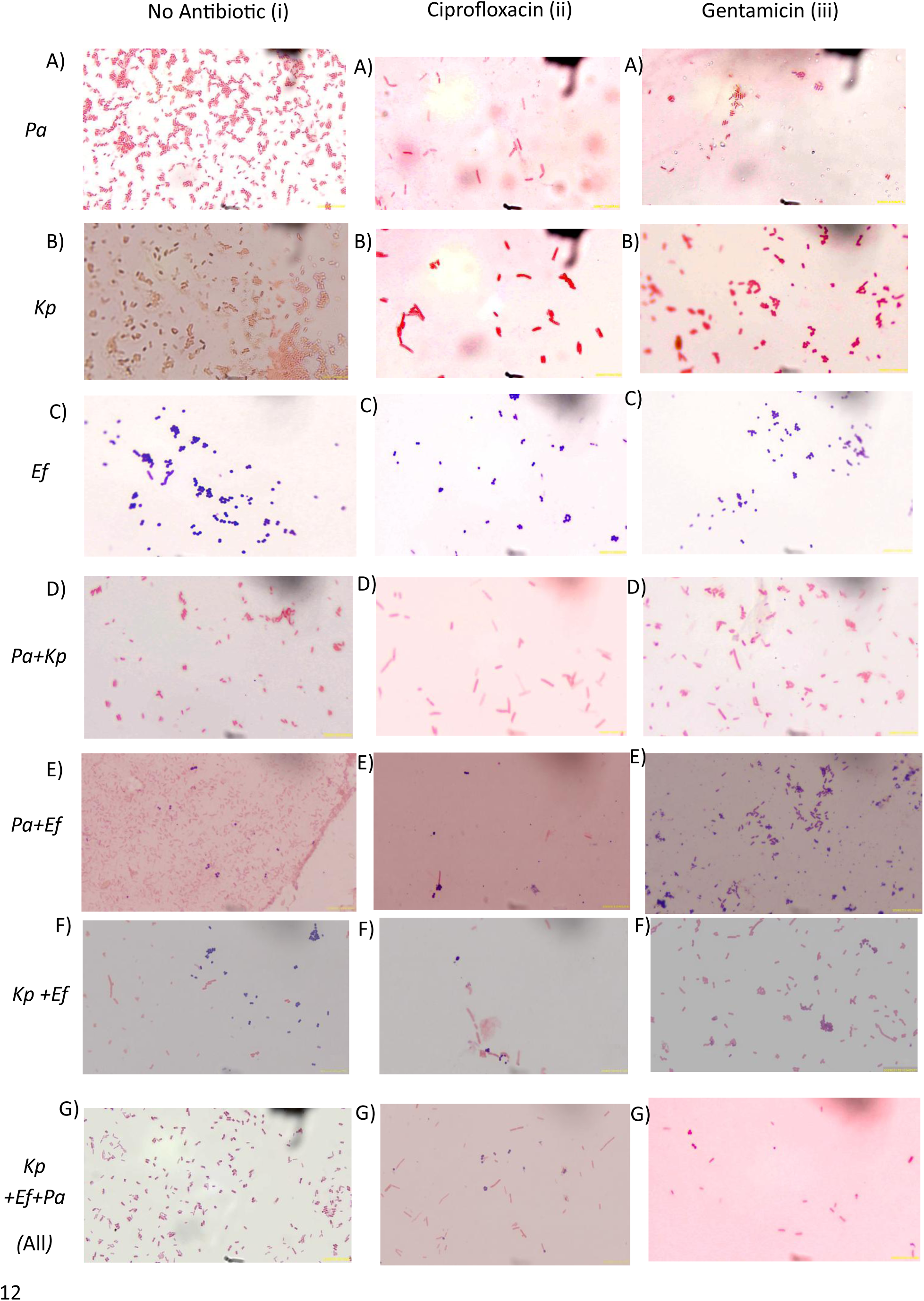
Adhesion assay for mono- and polymicrobial cultures under antibiotic treatments. Rows A–G represent isolate combinations; Columns i-iii represent untreated, ciprofloxacin, and gentamicin conditions respectively. Pa (A) and Kp (B) formed dense uniform monolayers under untreated condition; sub-MIC ciprofloxacin reduced adhesion and induced Kp filamentation, while gentamicin caused only moderate reduction. Ef (C) showed sparse adhesion with relative gentamicin tolerance. Pa+Kp (D) showed reduced co-culture adhesion versus monocultures, with severe and moderate clearing under ciprofloxacin and gentamicin respectively. Pa+Ef (E) demonstrated Pa-associated Ef retention, with distinct Ef microcolonies under gentamicin. Kp+Ef (F) showed patchy untreated colonisation with Ef becoming proportionally dominant under gentamicin pressure. Triple-species culture (G) formed dense untreated biofilms that were markedly reduced under ciprofloxacin, with residual elongated Kp filaments observed. Microscopy was done at 100x magnification and the scale is 10 µm.

### Growth Dynamics Under Flow Without Antibiotic Treatment

To establish the environmental impact on community assembly, we first compared growth dynamics in the absence of antibiotic stress. Log₂ fold change on the catheter and filter (i.e. dispersed cells) was calculated relative to the corresponding static control condition for each species and combination. Under antibiotic-free conditions, all three species showed increased biomass in monoculture compared with static controls (Figure 3a). Pa showed the greatest accumulation in both the catheter-associated (+17.38 log₂ fold change) and dispersed (+17.25) fractions. Ef showed substantial catheter-associated growth (+10.70) with a lower dispersed signal (+5.06), whereas Kp localized preferentially to the dispersed fraction (+5.49) over the catheter surface (+1.48). In dual-species communities, the Kp + Ef co-culture maintained high biomass in both fractions (Kp: +9.17 catheter, +8.97 dispersed; Ef: +10.55 catheter, +9.10 dispersed), consistent with cooperative growth under flow.

**Figure 3.**
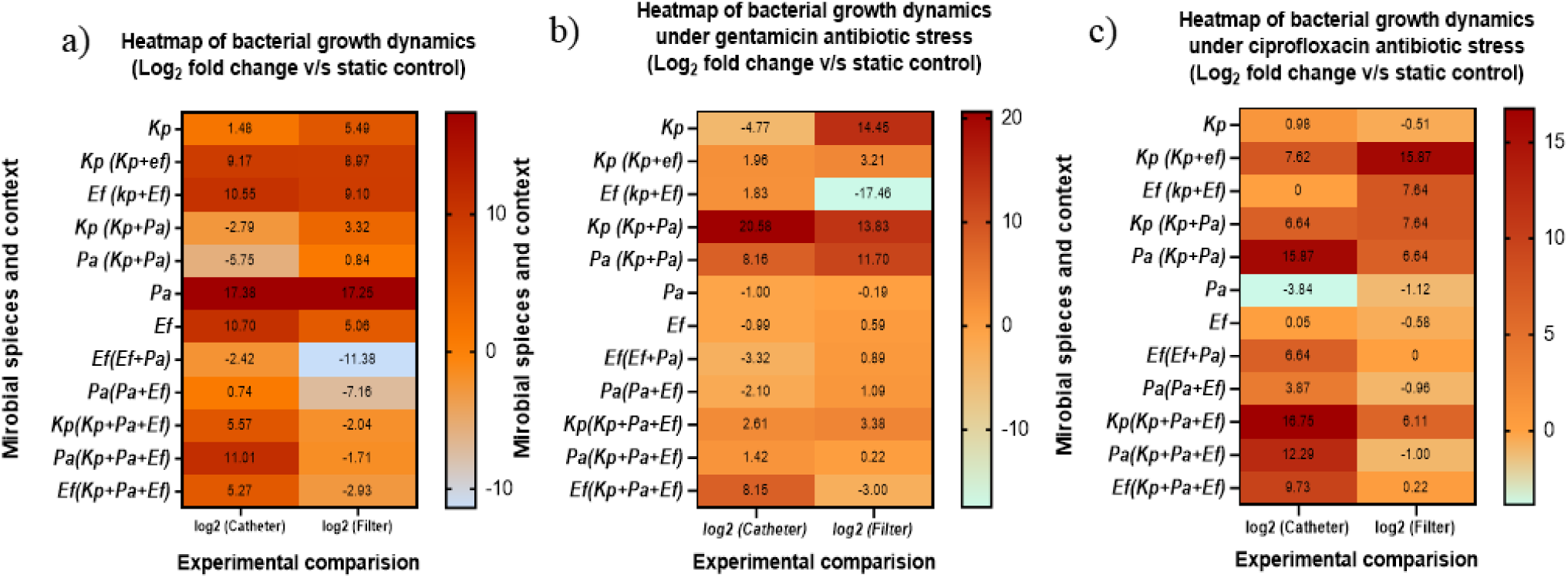
Heatmaps of log₂ fold changes in colony-forming units (CFU) of mono- and polymicrobial biofilm cultures under flow. Values represent changes in CFU counts relative to static baseline controls for *Klebsiella pneumoniae* (Kp), *Pseudomonas aeruginosa* (Pa), and *Enterococcus faecium* (Ef). Growth dynamics are tracked across catheter-associated (surface-attached) and filter (effluent/dispersed) populations. (a) Antibiotic-free control. Flow increased monoculture catheter biomass (Pa maximum: +17.38 log₂ FC). Kp+Ef grew cooperatively; Pa+Ef showed competitive inhibition (Pa effluent: -11.38 log₂ FC). Triple cultures showed a Kp-mediated buffering effect, recovering Pa catheter biomass (+11.01 log₂ FC). (b) Sub-MIC gentamicin. Kp monoculture showed maximum tolerance (+15.16 catheter; +14.45 filter log₂ FC). Kp+Pa displayed synergistic survival (Kp: +20.58; Pa: +8.16 log₂ FC catheter). Ef showed niche-specific protection in Kp+Ef (+1.83 catheter vs. -17.46 filter log₂ FC).(c) Sub-MIC ciprofloxacin. Pa monoculture was suppressed (catheter: -7.16; filter: -4.44 log₂ FC) but recovered in Pa+Ef (+6.64 log₂ FC catheter). Kp showed enhanced effluent dispersal in Kp+Ef (+15.87 log₂ FC) and dominated triple-species catheter populations (+16.75 log₂ FC).

On the contrary, Pa+Kp co-culture showed mutual suppression of surface colonization with reduced catheter abundance (Pa: −5.75; Kp: −2.79) and weak dispersal (Pa: +0.84; Kp: +3.32). Similar trends of competitive inhibition were observed in Pa+Ef co-culture, as Pa was suppressed across both fractions (catheter: −2.42; dispersed: −11.38), while Ef surface accumulation remained limited (+0.74) alongside reduced dispersal (−7.16). In triple-species communities, Pa catheter abundance dramatically increased (+11.01) compared to the Pa+Ef co-culture, whereas the dispersed fraction did not differ significantly (−1.71). Kp and Ef maintained moderate catheter growth (+5.57 and +5.27, respectively) but decreased dispersal ability (−2.04 and −2.93), indicating that in triple community integration restricts overall dispersal.

### Growth Dynamics Under Flow in presence of Gentamicin

Under flow conditions with gentamicin, species survival and interspecies interactions were differentially affected across catheter-associated and dispersed fraction (Figure 3b). In monoculture, Kp exhibited strong susceptibility in the catheter-associated fraction (−4.77), contrasting with high abundance in the dispersed fraction (+14.45), indicating that fluid flow increases susceptibility and promotes biofilm detachment and clearance from the catheter surface under gentamicin exposure. Pa showed minimal growth in both fractions (−1.00 catheter; −0.19 dispersed), while Ef showed a comparatively modest decline in the catheter fraction (−0.99), while its dispersed fraction increased slightly (+0.59). In dual-species communities, Kp maintained positive growth when paired with Ef (+1.96 catheter; +3.21 dispersed), though at reduced levels relative to monoculture. Ef displayed a niche-specific response in the Kp + Ef co-culture: modest catheter-associated growth (+1.83) contrasted with marked suppression in the dispersed fraction (−17.46), indicating catheter-associated protection. The Kp + Pa co-culture showed enhanced growth for both species in both fractions (Kp: +20.58 catheter, +13.83 dispersed; Pa: +8.16 catheter, +11.70 dispersed), consistent with synergistic survival under gentamicin stress. In contrast, the Pa + Ef co-culture showed limited or negative growth for Pa (−2.10 catheter; +1.09 dispersed) Ef (−3.32/+0.89), consistent with competitive suppression. In triple-species communities, all species showed moderate catheter-associated growth (Kp: +2.61; Pa: +1.42; Ef: +8.15), while dispersed-fraction abundance was reduced or variable (Kp: +3.38; Pa: +0.22; Ef: −3.00). Catheter-associated fractions consistently showed higher or more stable growth than dispersed fractions across conditions.

### Growth Dynamics Under Flow in presence of Ciprofloxacin

Under flow conditions, biomass was redistributed across catheter-associated and dispersed fractions rather than uniformly suppressed (Figure 3c). In monoculture, Pa was inhibited in both the catheter-associated (−3.84 log₂ fold change) and dispersed (−1.12) fractions. Kp demonstrated a minor positive growth in in catheter (+0.98) with reduction in disperesed fraction (−0.51), indicating survival without a proliferative advantage. Ef showed negligible catheter-associated change (+0.05) with marginal suppression in the dispersed fraction (−0.58). In dual-species communities, the Pa + Ef co-culture showed strong recovery of Pa and Ef in the catheter-associated fraction (Pa: +3.87; Ef: 6.64) relative to respective monocultures, consistent with a protective interaction, while the dispersed fraction of both organisms was minimal (Pa: -0.96; Ef: 0). The Pa + Kp co-culture showed further enhancement of catheter-associated abundance for both species (Pa: +15.87; Kp +6.64) along with dispersal (Pa: +6.64; Kp +7.64), consistent with synergistic growth under antibiotic stress. In the Kp + Ef co-culture, Kp abundance increased markedly in the dispersed fraction (+15.87) relative to monoculture, indicating ciprofloxacin-associated dispersal in the presence of Ef. In triple-species communities, Kp dominated catheter-associated populations (+16.75), exceeding both Pa (+12.29) and Ef (+9.73), indicating competitive dominance under polymicrobial conditions. Catheter-associated fractions consistently showed higher log₂ fold changes than dispersed fractions across all community compositions, consistent with enhanced ciprofloxacin tolerance in surface-associated populations.

### Antibiotic survival under flow and static conditions

To determine how flow alters drug tolerance, percentage survival was calculated by comparing treated CFU against untreated controls under both flow and static conditions. Quantitate evaluation of biomass was further supported by absolute colony-forming unit (CFU) measurements, which confirmed that flow increased baseline CFU for all three species in monoculture relative to static controls (Table 1). This effect was most pronounced in Pa, which reached 5.44 × 10¹³ CFU mL⁻¹ under flow compared to 3.20 × 10⁸ CFU mL⁻¹ under static conditions. Notably, CFU quantification revealed that increased biomass under flow conditions was not associated with enhanced antibiotic survival. Survival of Kp in the presence of gentamicin was very low (0.03%) under static conditions, and decreased further when in the flow condition (0.00% corresponding to 6.00 × 10³ CFU mL⁻¹). Pa, despite its substantially higher biomass under flow, survival fell sharply, from 15.00% to 4.41 × 10⁻⁵% in presence of gentamicin. In contrast, Ef demonstrated a distinct phenotypic response under static conditions, where sub-MIC gentamicin exposure elicited a hormetic effect, increasing the population density to 353% relative to the untreated control. This survival advantage was entirely suppressed under flow conditions, where Ef viability declined to 0.11%. Parallel to the trends seen with gentamicin, overall viability trends showed that Pa, despite its substantially higher biomass under flow, experienced a sharp decline in survival from 0.06% to 2.57 × 10⁻⁸% in presence of ciprofloxacin. A comparable reduction was observed for Ef in presence of ciprofloxacin exposure (20.00% static versus 0.01% flow). Kp survival in presence of ciprofloxacin was similarly low, falling to 0.04% under static conditions and 0.03% under flow. Across all species and antibiotic conditions, flow promoted biofilm accumulation in monoculture while reducing antibiotic survival relative to static conditions.

**Table 1.** Baseline biofilm density and survival of monocultures under flow and static conditions.

| Species | Condition | Control<br>CFU/mL | Gentamicin<br>CFU/mL | % Survival<br>(Gent) | Ciprofloxacin<br>CFU/mL | % Survival<br>(Cipro) |
| --- | --- | --- | --- | --- | --- | --- |
| *Kp* | Static | $4.80 \times 10^8$ | $1.64 \times 10^5$ | 0.03% | $2.00 \times 10^5$ | 0.04% |
| | Flow<br>(Cath) | $1.34 \times 10^9$ | $6.00 \times 10^3$ | 0.00% | $3.94 \times 10^5$ | 0.03% |
| *Pa* | Static | $3.20 \times 10^8$ | $4.80 \times 10^7$ | 15.00% | $2.00 \times 10^5$ | 0.06% |
| | Flow<br>(Cath) | $5.44 \times 10^1$<br>3 | $2.40 \times 10^7$ | $4.41 \times 10^{-5}$<br>% | $1.40 \times 10^4$ | $2.57 \times 10^{-8}$<br>% |
| *Ef* | Static | $3.00 \times 10^6$ | $1.06 \times 10^7$ | 353%* | $6.00 \times 10^5$ | 20.00% |
| | Flow<br>(Cath) | $5.00 \times 10^9$ | $5.34 \times 10^6$ | 0.11% | $6.20 \times 10^5$ | 0.01% |
| *Values >100% indicate hormetic growth induction at sub-MIC levels. |  |  |  |  |  |  |

### Relative abundance and species dominance in co-culture

#### In the absence of antibiotic exposure

Flow altered species dominance and community composition relative to static conditions across all pairings (Table 2a). In the Kp + Ef co-culture, Kp remained dominant under both conditions but its relative abundance decreased under flow (86.67% → 71.43%), with a corresponding increase in Ef (13.33% → 28.57%), consistent with reduced competitive exclusion. In the Pa + Ef co-culture, a dominance inversion was observed: Pa dominated under static conditions (84.21%), whereas Ef emerged as the abundant species under flow (62.50%). In the Kp + Pa co-culture, Pa dominated under static conditions (89.01%), while flow produced near-equal co-dominance (Kp: 49.15%; Pa: 50.85%). In the triple-species community, the dominant species changed between conditions: Kp was dominant under static conditions (59.43%), while flow produced a pronounced shift to Pa dominance (89.74%), with both Kp (7.69%) and Ef (2.56%) sharply reduced, indicating an abrupt flow-driven change in dominant species rather than a gradual redistribution.

**Table 2.** Relative abundance and species dominance in mixed species biofilms under static and flow conditions.

| Community | Environment | *Kp* (%) | *Pa* (%) | *Ef* (%) | Dominant Species |
| --- | --- | --- | --- | --- | --- |
| **Kp + Ef** | Static | 86.67 | 0.00 | 13.33 | **Kp** |
|  | Flow (Cath) | 71.43 | 0.00 | 28.57 | **Kp** |
| **Kp+Pa** | Static | 10.99 | 89.01 | 0.00 | **Pa** |
|  | Flow (Cath) | 49.15 | 50.85 | 0.00 | **Pa+Kp (equal)** |
| **Pa+Ef** | Static | 0.00 | 84.21 | 15.79 | **Pa** |
|  | Flow (Cath) | 0.00 | 37.50 | 62.50 | **Ef** |
| **Triple** | static | 59.43 | 16.04 | 24.53 | **Kp** |
|  | Flow (Cath) | 7.69 | 89.74 | 2.56 | **Pa** |

| Community | Environment | *Kp* (%) | *Pa* (%) | *Ef* (%) | Dominant Species |
| --- | --- | --- | --- | --- | --- |
| **Kp + Ef** | Static | 75.00 | 0.00 | 25.00 | **Kp** |
|  | Flow (Cath) | 76.64 | 0.00 | 23.36 | **Kp** |
| **Kp+Pa** | Static | 82.05 | 17.95 | 0.00 | **Kp** |
|  | Flow (Cath) | 100.00 | 0.00 | 0.00 | **Kp** |
| **Pa+Ef** | Static | 0.00 | 12.42 | 87.58 | **Ef** |
|  | Flow (Cath) | 0.00 | 5.71 | 94.29 | **Ef** |
| **Triple** | **Static** | **58.97** | **30.77** | **10.26** | **Kp** |
|  | **Flow (Cath)** | **10.67** | **2.44** | **86.89** | **Ef** |

C)Percentage relative abundance data for Kp, Pa, and Ef dual- and triple-species combinations under ciprofloxacin treatment
| **Community** | **Environment** | ***Kp* (%)** | ***Pa* (%)** | ***Ef* (%)** | **Dominant Species** |
| --- | --- | --- | --- | --- | --- |
| **Kp + Ef** | **Static** | **0.33** | **0.00** | **99.67** | **Ef** |
|  | **Flow (Cath)** | **38.82** | **0.00** | **61.18** | **Ef** |
| **Kp + Pa** | **Static** | **50.00** | **50.00** | **0.00** | **Pa+Kp (equal)** |
|  | **Flow (Cath)** | **98.28** | **1.72** | **0.00** | **Kp** |
| **Pa + Ef** | **Static** | **0.00** | **2.38** | **97.62** | **Ef** |
|  | **Flow (Cath)** | **0.00** | **14.29** | **85.71** | **Ef** |
| **Triple** | **Static** | **0.38** | **0.76** | **98.85** | **Ef** |
|  | **Flow (Cath)** | **32.35** | **2.94** | **64.71** | **Ef** |

#### In the presence of gentamicin

(Table 2b) For the Kp + Ef co-culture, Kp was dominant under both static (75.00%) and flow (76.64%) conditions, with Ef a consistent minority (25.00% static; 23.36% flow), indicating stable Kp dominance. In the Pa + Ef co-culture, Ef was strongly dominant under both static (87.58%) and flow (94.29%) conditions. In the Kp + Pa co-culture, Kp increased from 82.05% under static conditions to 99.996% under flow, whereas Pa declined from 17.95% to 0.004%. In the triple-species community, Kp was dominant under static conditions (Kp: 58.97%; Pa: 30.77%; Ef: 10.26%); under flow, Ef became the most abundant species (86.89%), with Kp and Pa decreasing to 10.67% and 2.44%, respectively.

#### In the presence of ciprofloxacin

(Table 2c) the Kp + Ef co-culture, Ef was near-exclusively dominant under static conditions (99.67%), and remained dominant under flow (Kp: 38.82%; Ef: 61.18%). In the Pa + Ef co-culture, Ef maintained its dominance across both conditions (97.62% static; 85.71% flow). In the Kp + Pa co-culture, co-dominance under static conditions (Kp: 50.00%; Pa: 50.00%) shifted to near-exclusive Kp dominance under flow (Kp: 98.28%; Pa: 1.72%), indicating that continuous flow altered competitive outcome independently of antibiotic exposure. In the triple-species community, Ef dominated under both static (98.85%) and flow (64.71%) conditions, with Pa reduced to a minor presence (0.76% static; 2.94% flow) and Kp intermediate under flow (32.35%); this indicates strong Ef dominance rather than balanced polymicrobial biofilm formation in response to ciprofloxacin exposure.

### Live/dead staining of biofilm communities under flow and static conditions

#### In the absence of any antibiotics

flow resulted in pronounced increase in live cell counts across species and community compositions (Figure 4,5). For monoculture, Pa (Figure 4a-c) and Kp (Figure 4g-i) exhibited increases in total biomass and live cell counts under flow relative to static conditions, while there was a decrease in viable counts for Ef (Figure 4d-f). Kp displayed a morphological transition from clustered, aggregated growth under static conditions to dispersed, uniform growth under flow, accompanied by the emergence of filamentous structures, suggesting early stage filamentation (Figure 4g-h). In co-culture, flow increased live cell counts across all combination relative to static conditions (Figure 5). In the Pa + Kp co-culture (Figure 5g-i), rod-shaped cells appeared reduced in size under flow, indicating cellular remodelling due to stress.

**Figure 4.**
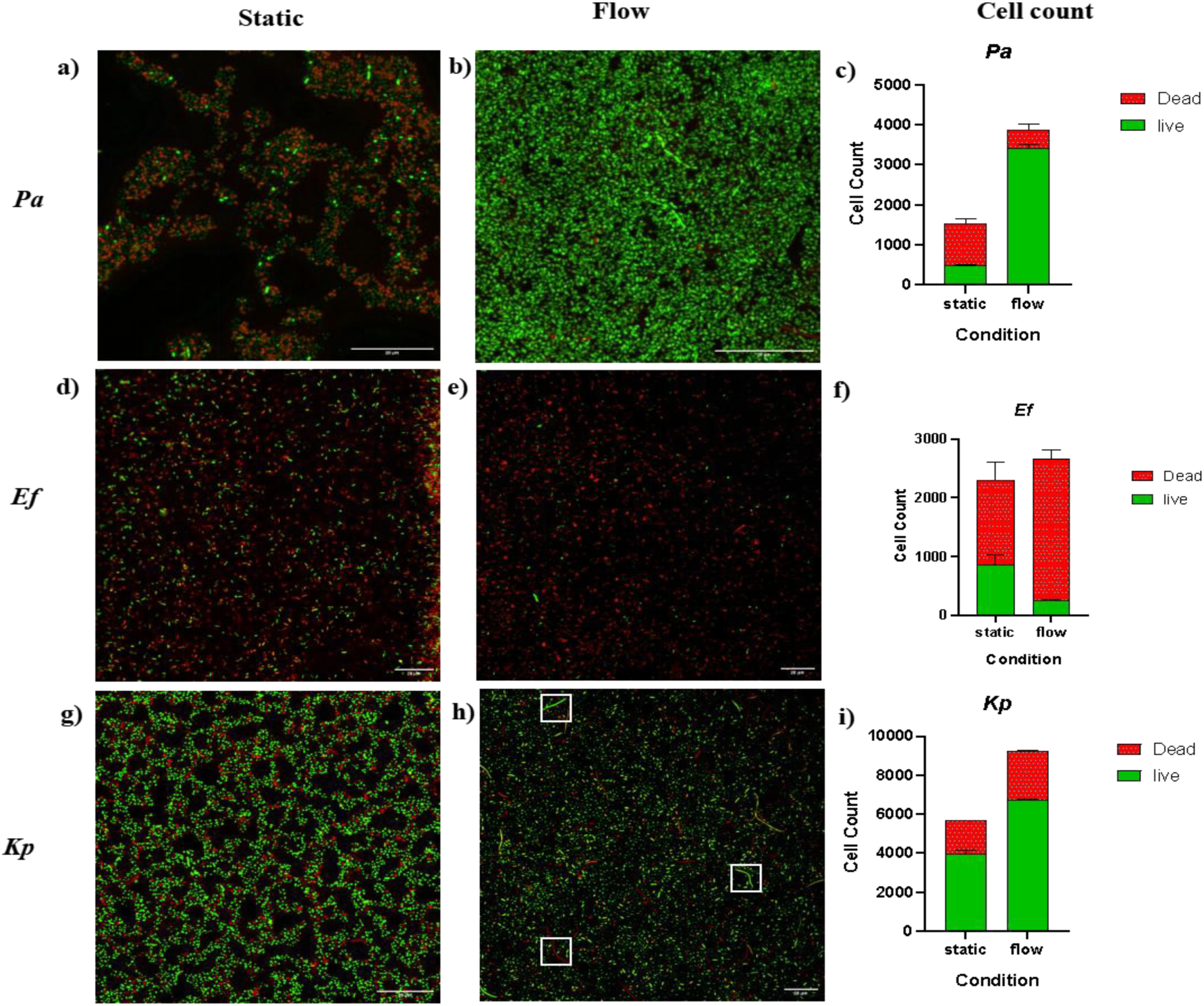
Confocal laser scanning microscopy (CLSM) visualization and cell count quantification of monoculture biofilms under antibiotic-free conditions. *P. aeruginosa* (Pa) and *K. pneumoniae* (Kp) biofilms exhibited substantial increases in total biomass and live cell counts under flow conditions (b, h respectively) relative to static conditions (a, g respectively), with Kp displaying a morphological transition from clustered aggregates to uniform, filamentous structures under flow (h). Conversely, *E. faecium* (Ef) showed a reduction in viable cell numbers under flow (e) compared to static conditions (d). Graphical representations of live and dead cell count for each isolate are shown in (c, f, i). Scale is 10µm.

**Figure 5.**
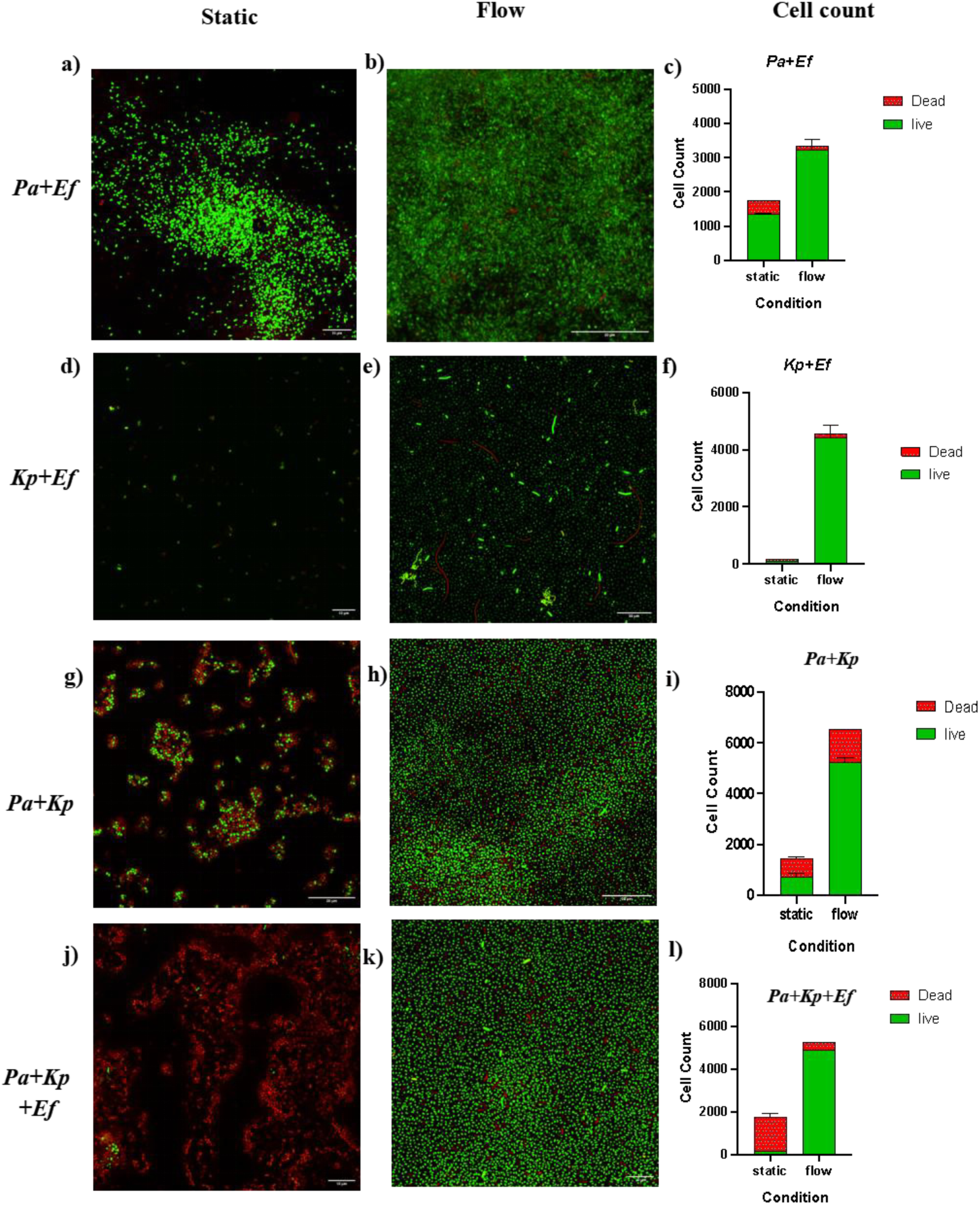
Confocal laser scanning microscopy (CLSM) visualization and cell count quantification of polymicrobial biofilms under antibiotic-free conditions. Flow conditions markedly increased live cell counts across all dual-species pairings (b, e, h) and the triple-species community (k) relative to their respective static controls (a, d, g, j). In the Pa+Kp dual-species culture under flow (h), rod-shaped cells displayed a visible reduction in size compared to static conditions (g). Graphical representations of live and dead cell count for each community layer are shown in (c, f, i, l). Scale as indicated directly on the individual micrographs.

#### Gentamicin exposure

resulted in significant variations in bacterial viability and clustering pattern between static vs. flow cultures (Figure 6,7). In monocultures, static Pa biofilms showed reduced biomass with high viability, whereas flow induced higher biomass with low viability cells indicating that flow enhanced susceptibility (Figure 6a-c). As opposed; Ef under static conditions resulted in limited viability, while flow promoted both increased cell numbers and a higher proportion of live cells, accompanied by chain-like morphology indicative of active growth and division under flow (Figure 6d-f). Static Kp culture formed dense, viable clusters that disrupted under flow into dispersed colony with high non-viable cell (Figure 6g-i). In co-culture the Pa+Ef co-culture resulted in superior viability under static conditions, but shifted to an Ef-dominated viability under flow (Figure 7a-c). The Kp+Ef co-culture exhibited increased cell death and rod shortening under flow (Figure 7d-f). The Pa+Kp co-culture also demonstrated increased susceptibility under flow (Figure 7g-h). In triple-species biofilms, flow reduced overall viability (Figure 7j-l); however, Ef formed localized aggregates surrounding rod-shaped cells, indicating antibiotic stress-induced spatial reorganization (Figure 7k).

**Figure 6.**
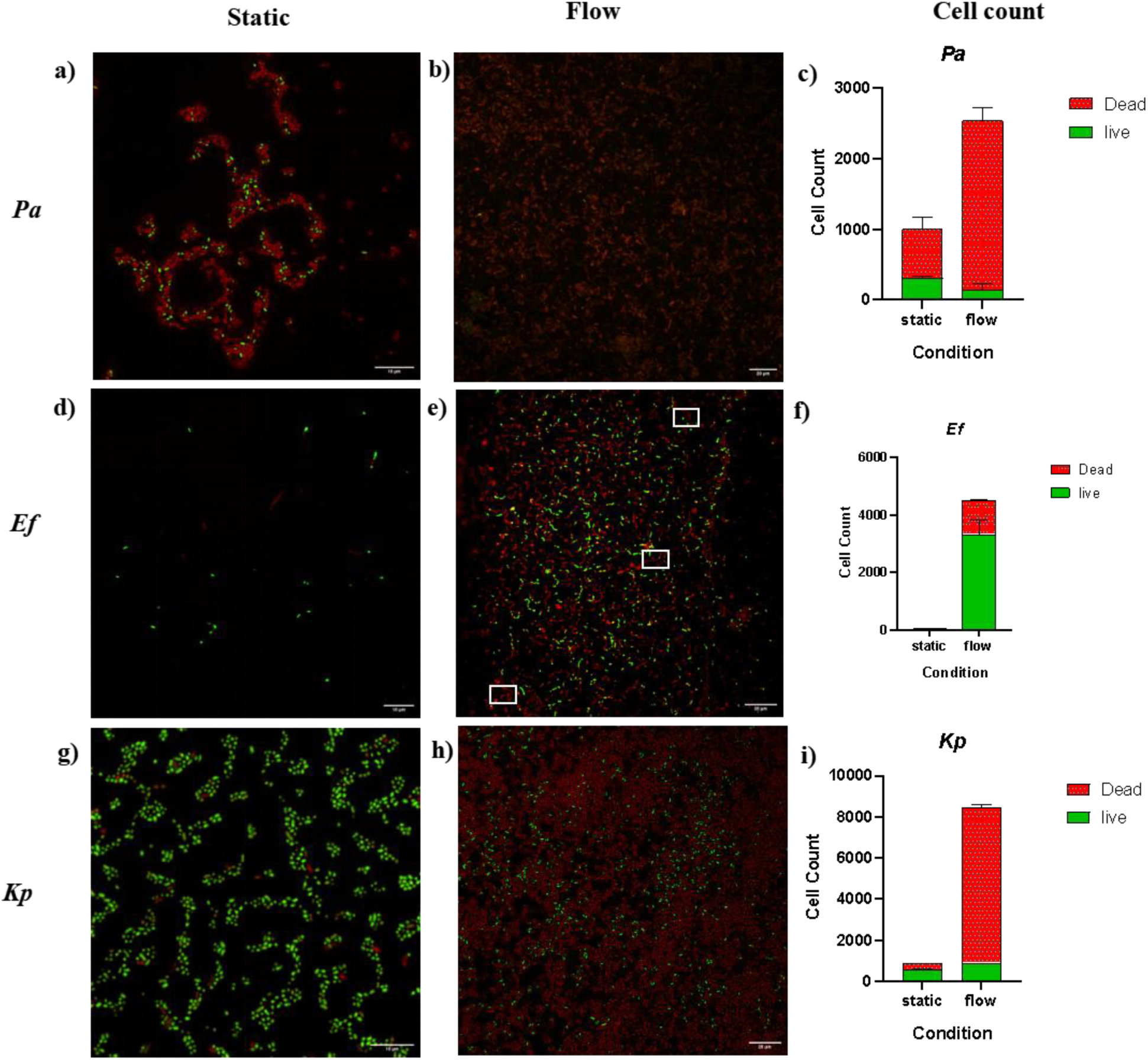
Confocal laser scanning microscopy (CLSM) visualization and cell count quantification of monoculture biofilms under subinhibitory gentamicin stress. Static Pa biofilms sustained a lower total cell density with higher relative viability (a), whereas flow increased total biomass dominated by non-viable cells (b). Ef under static conditions resulted in limited viability (d), while flow conditions promoted increased total cell numbers, a higher proportion of live cells, and a chain-like growth morphology (e). Static Kp formed dense, viable cell clusters (g) that disrupted under flow into dispersed populations with high non-viable counts (h). Graphical representations of live and dead cell count for each isolate are shown in (c, f, i). Scale bars are indicated directly on the individual micrographs.

**Figure 7.**
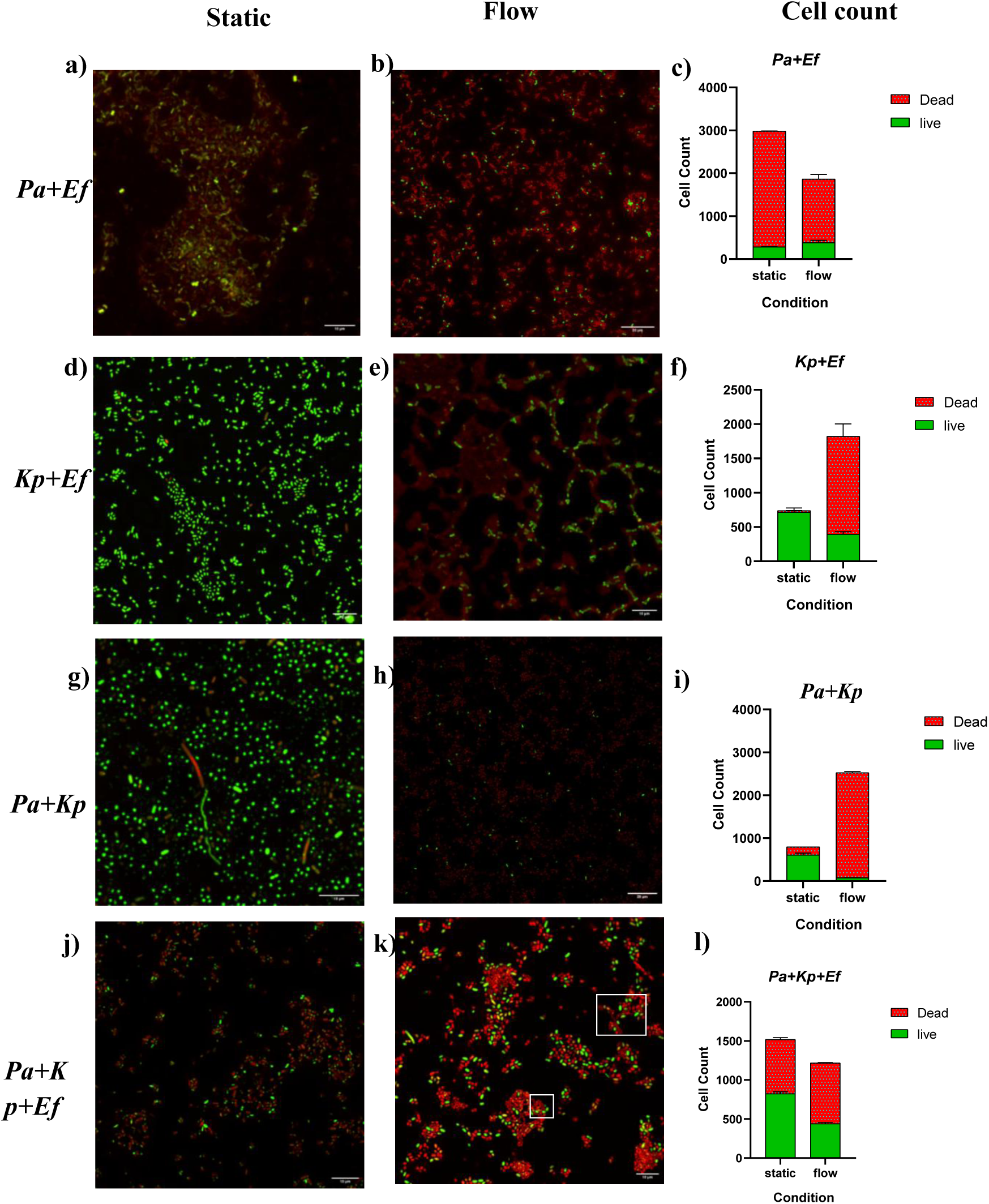
Confocal laser scanning microscopy (CLSM) visualization and cell count quantification of polymicrobial biofilms under subinhibitory gentamicin stress. Pa+Ef maintained superior overall viability under static conditions (a) but shifted to an Ef-dominated live cell population under flow (b). Kp+Ef exhibited increased cell death and rod shortening under flow conditions (e) relative to static controls (d). In triple-species biofilms, flow reduced overall viability (k), with Ef forming localized aggregates surrounding rod-shaped cells compared to the spatial distribution observed under static conditions (j). Graphical representations of live and dead cell count for each community layer are shown in (c, f, i, l). Scale bars are indicated directly on the individual micrographs.

#### Ciprofloxacin

induced the largest reduction in total cell number among all experimental conditions tested (Figure 8,9). Filamentation was the prevalent morphology observed in response to ciprofloxacin across conditions, particularly evident in Kp and predominantly non-viable in Pa. In monoculture, viability of Pa cultures under static conditions was greater, and flow resulted in reduced survival and mainly non-viable filamentous formation (Figure 8a-c). Ef exhibited increased live cell counts under flow relative to static conditions, demonstrating relative ciprofloxacin tolerance under shear (Figure 8d-f). Kp exhibited higher rates of survival in flow compared to the static condition, with significant filamentation being observed, suggestive of the inhibitory effect of ciprofloxacin on cell division (Figure 8g-i). In co-culture, ciprofloxacin significantly decreased overall biomass in all pairings tested (Figure 9). In Pa + Ef co-culture, survival in flow was limited only to cells from Ef, indicating a selective survival advantage under concurrent antibiotic and flow stress (Figure 9a-b). In the Kp + Ef, static conditions showed spatially organized mixed viability with Ef localizing at the termini of Kp filaments (Figure 9d); under flow, this shifted toward increased cell death and extensive filamentation (Figure 9e-f). In the Pa + Kp co-culture, static conditions supported higher viability with evidence of early filamentation, whereas flow promoted extensive filament formation that was predominantly non-viable, accompanied by reduced live cell counts (Figure 9g-i). In the triple-species community, total cell numbers declined substantially; however, under flow, live cell counts were relatively preserved alongside reduced dead cell detection (Figure 9j-l), consistent with either selective removal of non-viable biomass or persistence of a ciprofloxacin-tolerant subpopulation.

**Figure 8.**
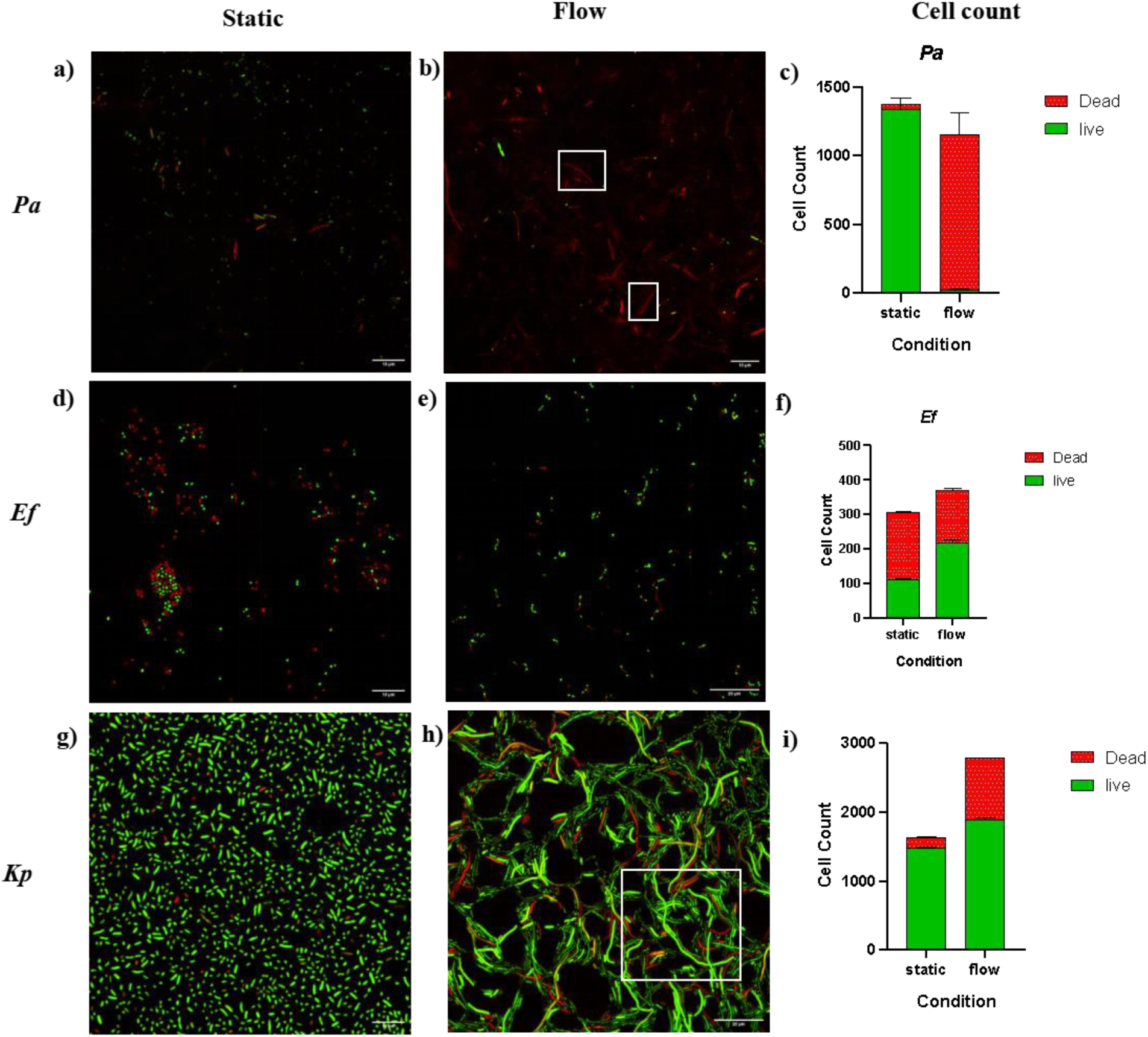
Confocal laser scanning microscopy (CLSM) visualization and cell count quantification of monoculture biofilms under subinhibitory ciprofloxacin stress. Pa viability was preserved under static conditions (a), whereas flow reduced overall survival and was accompanied by predominantly non-viable filamentous structures (b). Ef exhibited increased live cell counts under flow (e) relative to static conditions (d). Kp showed enhanced survival under flow compared to static controls (g), coupled with pronounced filamentation (h) consistent with ciprofloxacin-induced inhibition of cell division. Graphical representations of live and dead cell count for each isolate are shown in (c, f, i). Scale bars are indicated directly on the individual micrographs.

**Figure 9.**
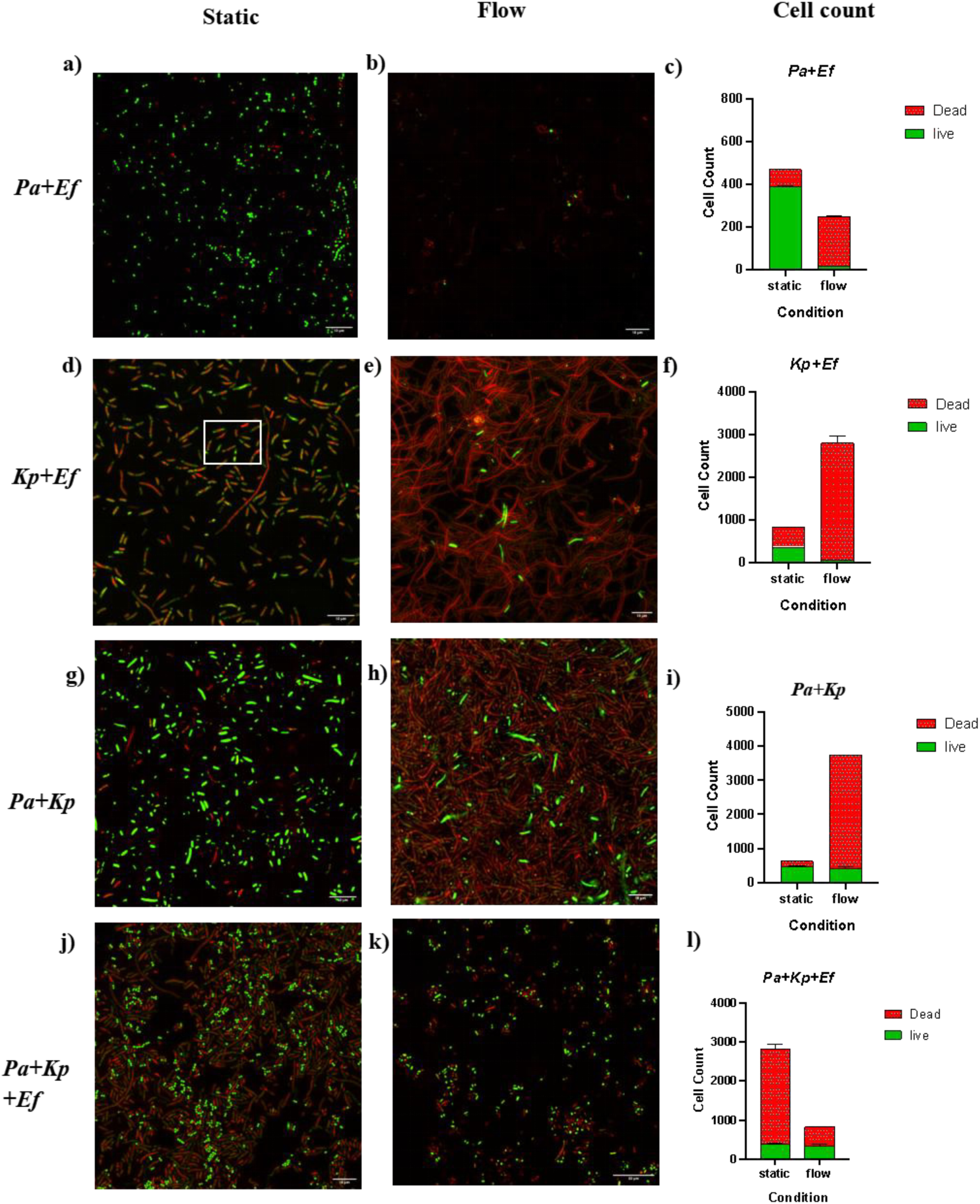
Confocal laser scanning microscopy (CLSM) visualization and cell count quantification of polymicrobial biofilms under subinhibitory ciprofloxacin stress. In Pa+Ef, viable cells under flow (b) were largely restricted to Ef compared to static conditions (a). In Kp+Ef, static conditions showed mixed viability with Ef localizing at the termini of Kp filaments (d) increased cell death and extensive filamentation observed under flow (e). In Pa+Kp, static showed higher viability with early filamentation (g), whereas flow had predominantly non-viable filament formation accompanied by reduced live cell counts (h). In the triple-species community, total cell numbers declined substantially, but live cell counts were relatively preserved under flow (k) relative to static conditions (j). Graphical representations of live and dead cell count for each community layer are shown in (c, f, i, l). Scale bars are indicated directly on the individual micrographs

## Discussion

This study tested whether continuous flow and sub-inhibitory antibiotic exposure jointly reshape competitive outcomes in polymicrobial catheter biofilms, a question that static in vitro models are unable to answer. Our results show that flow and antibiotic exposure together determine which species dominates a catheter community, and that this outcome is quite different from what static culture predicts. The clearest demonstration of this is *Enterococcus faecium*, an organism that is uniformly classified as a weak biofilm producer in monoculture assays, yet rose to community dominance under flow in co-culture with *Pseudomonas aeruginosa* (15.8% to 62.5% relative abundance). A second line of evidence points the same way: under sub-MIC ciprofloxacin, Ef microcolonies localised specifically at the tips of *K. pneumoniae*’s antibiotic-induced filaments, suggesting flow and antibiotic stress open more than one route to Ef persistence in a mature CAUTI biofilm not detectable by static models.

### Flow altered growth, survival, and dominance

Flow increased biomass for all three species in monoculture consistent with studies showing that flow enhances bacterial growth and viability by continuously replenishing nutrients and sustaining higher live-cell populations relative to static conditions (21, 30). Pa showed the largest gain (+17.38 catheter/+17.25 dispersed; CFU rising from 3.20 × 10⁸ to 5.44 × 10¹³ mL⁻¹). This may be attributed to shear-responsive surface sensing, which has been shown to elevate c-di-GMP and promote surface commitment in *P. aeruginosa* (31). Ef dominance in the catheter fraction (+10.70) aligns with established models of pilus-mediated adhesion in catheter-associated *Enterococcus* infections (32). Whereas, Kp’s shift toward the dispersed fraction (+1.48 catheter/+5.49 dispersed) mirrors the ’dispersal-favoring phenotypes’ described in recent literature, where biofilm-dispersed *K. pneumoniae* cells exhibit enhanced colonization efficiency compared to sessile cells (33). These findings indicate that community restructuring under flow is driven more by active adaptive strategies than by baseline adhesion alone.

Co-culture reversed the monoculture growth patterns. The Kp+Ef co-culture grew cooperatively in both fractions (Kp +9.17 catheter/+8.97 dispersed; Ef +10.55/+9.10), a finding that aligns with recent evidence showing *K. pneumoniae* metabolites can stimulate growth in catheter models in related Enterococcal species *E. faecalis* (34). While *E. faecalis* has been reported to inhibit *K. pneumoniae* in static, glucose-rich conditions (35), these results suggest that flow conditions may dilute inhibitory metabolites, allowing cooperative interactions to dominate. Conversely, both Pa co-cultures were suppressed. Pa+Kp showed mutual suppression (Pa −5.75, Kp −2.79), consistent with reports that Pa phenazines suppress *K. pneumoniae* (36), which, under flow, likely compounds with resource competition. Despite the competitive suppression observed in the Pa+Ef co-culture (Pa: −2.42 catheter/ −11.38 dispersed; Ef: +0.74/−7.16), flow inverted the community hierarchy, shifting Pa from 84.21% (static) to 37.50% relative abundance, while Ef rose to 62.5%. This indicates that flow does not necessarily favor Ef growth, but rather suppresses Pa’s retention more severely, effectively forcing a dominance inversion. This aligns with the ’niche-vacancy’ mechanism described by Holt et al. (2024), where *P. aeruginosa*’s intrinsic, periodic dispersal under flow actively vacates surface area, thereby creating spatial opportunities for co-inhabitants to persist (37).The Kp+Pa co-culture showed a related shift, moving from Pa dominance (89.01% static) to near-equal co-dominance (50.85% Pa, 49.15% Kp) under flow, and Kp+Ef stayed Kp-dominant throughout but fell from 86.67% to 71.43% as Ef increased.

In the triple-species community, Pa catheter growth recovered sharply (+11.01), suggesting that *K. pneumoniae* minimizes the *Pa–Ef* antagonism. This is consistent with a ’higher-order interaction’ phenomenon which is increasingly recognized as a primary driver of stability in complex microbial consortia where the pairwise competition between two species is fundamentally altered by the presence of a third (38) . While recent static, pairwise imaging shows that *P. aeruginosa* undergoes amensalism with *K. pneumoniae* and mutualism with *Enterococcus* on flat surfaces (39), our community dynamics behaved completely differently. Relative abundance shifted markedly rather than toward near-balance: Kp was dominant under static conditions (59.43%), and Pa became overwhelmingly dominant under flow (89.74%), while both Kp (7.69%) and Ef (2.56%) were sharply reduced. This realignment implies that baseline pairwise archetypes observed under static conditions may not accurately predict community behaviour under fluid shear stress in a flow environment. These results support the observation that polymicrobial outcomes in flow environments are driven by dynamic, species-specific interaction that cannot be fully predicted from pairwise data alone.

Live/dead staining aligned with these trends: flow increased live cell counts for almost every species and pairing, a result consistent with other catheter-relevant viability models (21). One notable exception was Ef, which declined under flow in monoculture, paralleling reports in the related species *E. faecalis*, where flow initiation caused a 3–4 log drop in adherent population within 4 h in catheter-calibrated drip-flow reactors — a phenotype absent in static assays of the same species (40). Whether *E. faecium* shares this vulnerability remains untested. Furthermore, Kp shifted from clustered static growth to dispersed growth with early filamentation under flow.

This is consistent with the broader understanding that flow-induced shear forces alter bacterial biofilm growth and morphology in gram-negative systems (41). Similarly, Pa+Kp rod cells appeared smaller under flow, this morphological change suggests a stress-induced structural remodelling unique to the co-culture environment under flow.

### Gentamicin reshaped survival and dominance under flow

In monoculture, Kp showed a divergent response to gentamicin: catheter-associated biomass declined (−4.77), while the dispersed fraction increased substantially (+14.45), suggesting gentamicin promoted detachment from the catheter surface rather than surface growth, consistent with Kp’s dispersal-favoring phenotype under flow (26). While sub-inhibitory gentamicin has been reported to promote *K. pneumoniae* growth and biofilm formation under static conditions (42), our observation under flow demonstrates net reduction on the catheter surface. Pa showed minimal growth (−1.00/−0.19); while literature suggests sub-MIC gentamicin can induce biofilm formation in *P. aeruginosa* via increased eDNA and ROS (43), our data suggests that the continuous shear and dilution effects of the flow environment likely override these inductive signals, shifting the balance toward growth suppression. Ef showed minor surface reduction in the catheter fraction (− 0.99); while sub-inhibitory concentrations of antibiotics are widely reported to induce biofilm formation in *Enterococcus* species (44) ,our findings are consistent with specific models— including *E. faecalis*—where gentamicin is shown to inhibit growth and biofilm development on the catheter surface (45).

In co-culture, Kp+Pa grew synergistically (Kp +20.58/+13.83; Pa +8.16/+11.70). While direct literature on this specific synergy under sub-MIC gentamicin is limited, our findings highlight a clear divergence from their typical competitive behaviour, pointing toward the need for further mechanistic investigation. In the Kp+Ef co-culture, Ef was favored within the catheter-associated biofilm (+1.83 catheter, −17.46 dispersed), which aligns with a potential mechanism where *K. pneumoniae*-derived factors stimulate the *Enterococcus* Fsr quorum-sensing system (34). In the Pa+Ef co-culture, both species were suppressed on the catheter surface (Pa −2.10; Ef −3.32), consistent with findings that *Enterococcus* species antagonize *P. aeruginosa* growth through metabolic byproducts like lactic acid and iron chelation (46). This inhibitory effect appears surface-specific, as antagonism did not persist in the dispersed fraction (Pa +1.09; Ef +0.89). Our observations suggest that this difference may arise because detachment moves cells into bulk flow, where these inhibitory metabolites are diluted, thereby reducing their localized impact. In the triple-species community, Ef showed the highest catheter growth (+8.15), indicating that the behaviour of each species depends on the entire polymicrobial community, not just pairwise interactions (47).

Survival data supported these trends. Kp survival plummeted under flow (0.00% from 0.03%), as did Pa (15.00% to 4.41 × 10⁻⁵%). Ef was the exception: sub-MIC gentamicin produced a hormetic effect under static conditions (353% of control), which was lost under flow (0.11% survival). Similar studies in *E. faecalis* have shown that aminoglycoside exposure upregulates the adhesion gene efaA (13, 13). *E. faecium* carries its own efaA homolog, efaAfm (48) which is a plausible, though unconfirmed, explanation here. The loss of this effect under flow suggests the mechanism relies on the local accumulation of a diffusible signal, which is disrupted by continuous shear stress.

Relative abundance reinforced this pattern. In the Kp+Ef co-culture, Kp remained dominant under both static (75.00%) and flow (76.64%) conditions. While Kp–*E. faecium* interactions remain understudied under continuous flow and antibiotic pressure, these observations run counter to a previous static study where *E. faecalis* suppressed Kp (35). However, it aligns with reports that Kp biofilm formation is not suppressed during *Enterococcus* co-inoculation under static baseline conditions (49). One possible contributing factor is that flow limits the local buildup of diffusible metabolic signals or secondary metabolites that might otherwise favor Ef in static settings, though this remains to be tested. The Pa+Ef co-culture remained Ef-dominant irrespective of flow (87.58%→94.29%) likely supported by the efaA -linked adhesion pathway as mentioned above (48), while Kp+Pa was already Kp-dominant under static conditions and became near-exclusively so under flow (82.05%→99.996%), mirroring the antibiotic-free pattern. The triple-species community was Kp-dominant, under static conditions (Kp 58.97%, Pa 30.77%, Ef 10.26%), shifting to near-exclusive Ef dominance under flow (86.89%). This indicates that Ef’s competitive advantage under flow is context-dependent, being most pronounced in the triple-species assemblage under combined effects of flow and antibiotic stress, where it displaces both Kp and Pa almost entirely. To our knowledge, this represents the first observation of flow-mediated *E. faecium* dominance within this tripartite community under sub-MIC gentamicin; we present this three-way competitive outcome as a novel observation of this system.

Live/dead staining matched these trends. Flow increased Pa biomass but reduced viability. This is consistent with reports that continuous-flow *P. aeruginosa* biofilms undergo localized cell death and lysis while maintaining a viable subpopulation (50) and that sub-MIC gentamicin independently stimulates matrix (alginate) production (51); the combined effect of flow and gentamicin together, as observed here, has not previously been reported. Ef showed the opposite trend: higher viability and a shift toward chain-like morphology under flow. This stress-adaptive pattern is plausible, given reports of antibiotic-induced restructuring (52) and gentamicin-induced biofilm upregulation (13) in *E. faecalis*. While these specific responses remain to be validated in *E. faecium* isolates, the current data strongly suggest a similar structural adaptation. In the triple-species community, Ef formed localized aggregates around rod-shaped cells under flow. This spatial arrangement has not been previously reported for this combination to our knowledge thus requiring further mechanistic investigation.

### Ciprofloxacin reshaped survival and dominance under flow

Ciprofloxacin elicited distinct responses across species. In monoculture, *P. aeruginosa* (Pa) remained vulnerable (−3.84/−1.12), consistent with reports that Pa biofilms require specific nutrient and shear conditions to build tolerance independent of biomass (41). Conversely, *K. pneumoniae* (Kp) showed little net change in either fraction (+0.98 catheter, −0.51 dispersed), indicating survival without a proliferative advantage, supporting evidence that Kp biofilms resist ciprofloxacin through limited drug penetration into the biofilm matrix (53). *E. faecium* (Ef) showed negligible catheter-associated change (+0.05). While sub-inhibitory ciprofloxacin has been reported to upregulate adhesion and biofilm related genes at the transcriptional level in *E. faecium* (54), this was not reflected in catheter-associated biomass under our flow conditions. In co-culture, Pa growth recovered significantly when paired with Ef (−3.84 → +6.64 catheter) or Kp (Pa +15.87; Kp +6.64), suggesting protective synergy. This parallels recent findings that *E. faecalis* antagonism enhances Pa antibiotic survival (46); whether this mechanism translates to *E. faecium* remains to be determined. Notably, the Kp+Ef co-culture increased Kp dispersal (+15.87 vs. −0.51 in monoculture), a novel observation suggesting Ef promotes Kp dispersal under fluoroquinolone stress. This is clinically concerning, as biofilm-dispersed *K. pneumoniae* are known to exhibit superior colonization and survival capabilities compared to planktonic cells (33), implying that incomplete antibiotic penetration may inadvertently promote dissemination rather than clearance (55).Finally, in the triple-species community, Kp dominated growth on catheter (+16.75) over Pa (+12.29) and Ef (+9.73), consistent with clinical surveys of recurring polymicrobial catheter biofilms (56).

Under flow conditions, viability plummeted for all monoculture. Pa’s viability fell sharply to 2.57×10^−8^%, likely reflecting the combined metabolic stress of flow and antibiotic exposure together, which increases susceptibility beyond either factor alone. Ef experienced a reduction from 20% to 0.01% under flow. While sub-inhibitory ciprofloxacin has been shown to upregulate adhesion genes in static *E. faecium* cultures (54), whether this response translates to a survival advantage under continuous flow remains untested. Finally, Kp exhibited smaller reduction in its viability, retaining a minor survivor fraction (0.03%) under ciprofloxacin, a finding consistent with reports that biofilm-forming *K. pneumoniae* in CAUTIs frequently harbor quinolone resistance (57).

In Pa+Ef co-culture, ciprofloxacin shifted relative abundance toward *E. faecium* dominance (97.62% static; 85.71% flow); while existing literature indicates that sub-inhibitory ciprofloxacin impairs *P. aeruginosa* quorum sensing, virulence, and biofilm formation in static or planktonic conditions (58, 59), our results demonstrate that this antibiotic-mediated competitive suppression persists, and intensifies, under continuous flow. In the Kp+Ef co-culture, ciprofloxacin drove near-exclusive Ef dominance under static conditions (99.67%), consistent with reports that *E. faecium* clinical isolates frequently carry multiple efflux pump determinants conferring high-level ciprofloxacin resistance (60). This dominance moderated under flow (Kp: 38.82%; Ef: 61.18%), consistent with evidence that continuous flow can disrupt the local antimicrobial-neutralizing gradients that protect resistant populations under static conditions, thereby increasing antibiotic effectiveness and reducing the competitive advantage such resistance confers (61). Conversely, Kp+Pa competition favored *K. pneumoniae* moving from co-dominance (50.0%) to near-exclusive Kp dominance (98.28%) upon the introduction of flow, mirroring the pattern of Kp dominance in nutrient-replete flow-through systems (62). This suggests that while antibiotics drive competitive shifts in the Pa+Ef co-culture, flow is the primary driver for the Kp+Pa shift, independent of antibiotic-induced competitive suppression. In the triple-species community, Ef remained dominant under both static (98.85%) and flow (64.71%) conditions, while Kp increased from a negligible static presence (0.38%) to a substantial minority under flow (32.35%); Pa remained marginal throughout (0.76% static; 2.94% flow). Kp’s gain here, despite a comparatively neutral monoculture response (+0.98 catheter), suggests that its competitive advantage under ciprofloxacin emerges specifically through polymicrobial interaction rather than intrinsic tolerance. Ef’s continued dominance likely reflects its higher baseline density and its own ciprofloxacin resistance (60).Pa’s minimal presence under both conditions is consistent with its ciprofloxacin sensitivity (41), potentially compounded by enterococcal antagonism (46).

Live/dead staining identified filamentation as the predominant morphological response to ciprofloxacin. Although SOS-mediated filamentation (63) is well-characterized, we observed these phenotypes under flow conditions. In Pa+Ef biofilms, survival was restricted almost entirely to *E. faecium*, mirroring antagonistic behaviors documented in other enterococci: specifically, recent reports have shown that *E. faecalis* antagonizes *P. aeruginosa* in mixed biofilms, reducing its viability to a small subpopulation (46). While the specific molecular mechanisms may differ between *Enterococcus* species, our data suggests that enterococcal antagonism remains a potent driver of community structure even under the shear stress of a flow model. Furthermore, Kp+Ef co-culture displayed *E. faecium* microcolonies localizing preferentially at the termini of *K. pneumoniae* filaments. Given that filamentation increases available attachment surface area, this spatial association, to our knowledge, has not been previously reported in CAUTI flow models thus representing a novel finding of this study.

### Why Standard Biofilm Models Fail to Predict Clinical Outcomes

In the absence of antibiotics, community composition alone determined which species dominated under flow, and this varied unpredictably across pairings i.e. no single organism dominated in every combination. Sub-inhibitory antibiotic exposure resolved this unpredictability into a consistent, antibiotic-specific pattern: gentamicin consistently drove Kp to dominance, while ciprofloxacin consistently drove Ef to dominance. This indicates that dominance in a polymicrobial catheter biofilm is not an intrinsic property of any single organism, but a function of which antibiotic the community happens to encounter.

Kp’s dominance under gentamicin was paradoxical. In monoculture, gentamicin nearly eliminated Kp where survival on the catheter surface fell to a negligible fraction of control. Yet in every polymicrobial pairing under the same drug, Kp emerged as the dominant species, moving toward near-total dominance in Kp+Pa. This is consistent with antibiotic-driven competitive release, whereby antibiotic-mediated suppression of competing species allows previously minor populations to expand despite no change in their intrinsic susceptibility (64).Given Kp’s already-recognized priority status, this finding shows that standard monoculture susceptibility testing, run under static conditions, can be actively misleading for Kp, since a drug that appears highly effective against it in isolation may instead simply clear its competitors under real flow conditions, leaving Kp to dominate once other species are removed from the picture.

Ef’s competitive dynamics were among the most striking findings of this study. Although *E. faecalis* is the standard laboratory model for CAUTI, *E. faecium* poses the greater clinical threat, carrying a wider array of antibiotic resistance determinants and driving most vancomycin-resistant enterococcal catheter infections (65, 66). Yet Ef remains understudied under flow, largely because the more genetically tractable *E. faecalis* has become the default biofilm model (67, 68). Our data shows that Ef’s dominance is an active, adaptive strategy. Flow appears to act on Ef through two independent routes. First, mechanically: Ef’s chain-like morphology under shear likely enhances surface retention and resistance to detachment, as demonstrated for *E. faecalis*, potentially contributing to its competitive advantage under flow (69). Second, through signalling: flow suppressed the Fsr quorum-sensing system of *E. faecalis* in a microfluidic endocarditis model, with biofilm growth and gentamicin tolerance rising only once bacteria were shielded from flow (70), suggesting flow actively restrains a repressive signal rather than merely failing to clear Ef. Ef also acquires antibiotic recalcitrance through polymicrobial facilitation from gram-negative partners (71). Together, these findings indicate that Ef’s persistence reflects community-driven behaviour rather than intrinsic resistance alone, and that effective management may require targeting its adaptation to polymicrobial, high-shear environments.

Several methodological limitations of this study should be acknowledged. This study employed a defined *in vitro* catheter flow model comprising three clinically relevant uropathogens, which permits controlled investigation of polymicrobial dynamics but it fails to capture ecological complexity of biofilms observed in clinical practice in the context of catheters. The 24-h experimental window captured early colonization and acute antibiotic adaptation; however, extended analyses beyond 48 h will be required to resolve persistence-associated community restructuring during biofilm maturation. Although CFU-based quantification robustly defined survival and competitive dynamics under flow, the molecular determinants underlying interspecies facilitation and flow-dependent protection were not directly resolved. Future genetic and transcriptional analyses will be required to establish mechanistic causality underlying these phenotypes.

## Conclusion

In conclusion, this study demonstrates that the reliance on static, single-species models fundamentally obscures the ecological realities of catheter-associated infections. In the absence of antibiotics, no single organism dominated across all community combinations under flow. Different species prevailed depending on which organisms were present together. Sub-inhibitory antibiotic exposure resolved this unpredictability into a consistent, antibiotic-specific pattern: gentamicin favored Kp, and ciprofloxacin favored Ef. This demonstrates that flow and antibiotic class together, not either alone, are active determinants of species dominance in polymicrobial catheter biofilms, and that the competitive fitness of both organisms is masked by standard laboratory assays. By demonstrating that treatment failure is often rooted in biofilm architectures and community behaviours invisible to conventional testing, this work exposes a significant disconnect in how current antibiotic strategies are calibrated. Ultimately, these results position flow as a critical ecological variable in CAUTI pathogenesis and argue that transitioning to dynamic, multi-species models is essential to effectively managing the complex, resilient biofilms encountered in the clinical environment.

## Materials and methods

### Bacterial strains and culture conditions

Clinical isolates of *Pseudomonas aeruginosa* (n=5), *Klebsiella pneumoniae* (n=5), and *Enterococcus facieum* (n=5) were recovered from urinary catheters of hospitalised patients at a tertiary care institution in Gujarat and Rajasthan, India. All strains were maintained as glycerol stocks and revived on Mueller Hinton agar (MHA) prior to each experiment. Routine culture was performed in Mueller-Hinton broth (MHB) or Luria-Bertani (LB) broth at 37°C with orbital shaking at 180 rpm. Experimental inocula were prepared by adjusting overnight cultures to an optical density of 0.08 OD₆₀₀ (approximately 10⁸ CFU mL⁻¹ equivalent to 0.5 McFarland standard) in MHB. For polymicrobial inocula, individual suspensions at 0.1 OD₆₀₀ were mixed 1:1 (v/v) for dual cultures or 1:1:1 (v/v/v) for triple cultures. Three representative isolates—*P. aeruginosa* J23, *K. pneumoniae* Top47, and *E. faecium* Ef50— were selected for biofilm flow experiments based on MIC profiles that provided comparable antimicrobial susceptibility distribution coverage within their respective species cohorts, avoiding extreme outlier values to maximize experimental comparability (supplementary Table 1, supplementary Figure 1)

### Biofilm quantification and categorisation

Biofilm formation across the full clinical isolate panel and representative mixed-species combinations was quantified using crystal violet (CV) microtiter plate assay as per the method by Stepanovic et al (72). 100 µL of cultures adjusted to OD₆₀₀ ≈ 0.08 were added to 50 µL MHB into sterile 96-well flat-bottom plates in triplicate and incubated at 37°C for 24 h. After removal of non-adherent cells by washing with sterile distilled water, adherent biofilm was fixed and stained with 150 µL of 0.1% crystal violet for 20 min, washed twice, and solubilised in 300 µL absolute ethanol for 30 min. Absorbance was read at 590nm. Isolates were classified as strong (OD > 4ODc), moderate (2ODc < OD ≤ 4ODc), or weak (ODc < OD ≤ 2ODc) biofilm producers, where ODc (cut-off OD) was defined as three standard deviations above the mean OD₅₉₀ of sterile MHB negative controls. Statistical analysis and graph were plotted using GraphPad Prism 8.0.

### Antimicrobial susceptibility testing and sub-MIC determination

Minimum inhibitory concentrations (MICs) of ciprofloxacin and gentamicin were determined by broth microdilution in MHB following Clinical and Laboratory Standards Institute (CLSI) guidelines, using serial two-fold dilution from the highest desired concentration. Bacterial inocula were prepared to 0.5 McFarland standard, diluted 1:100 in MHB. Plates were incubated at 37°C for 24 h. Growth was assessed by visual turbidity; MIC was defined as the lowest concentration producing no visible turbidity. Sub-MIC concentrations for monocultures were set at ½ × MIC, whereas for polymicrobial sub-MIC determination, the antibiotic concentration at which co-culture suspensions remained viable (i.e., permitted survival while not achieving complete inhibition) was used as the operational sub-MIC for subsequent flow experiments (supplementary Table 1).

### Cell adhesion assay

Surface-adhesion capacity of selected isolates was evaluated using a 6-well tissue culture plate-based adhesion assay. Bacterial cultures (OD₆₀₀ = 0.1 in MHB) were inoculated (5 ml) onto sterile glass coverslips (22 mm) placed in 6-well tissue culture plates. After 4 hours incubation at 37°C, coverslips were washed three times with 0.85% saline, heat-fixed, and subjected to Gram staining. Adherent cells were visualized at 100× magnification under oil immersion on a light microscope. Experiments were performed for all monoculture and co-culture combinations under antibiotic free, sub-MIC ciprofloxacin, and sub-MIC gentamicin conditions.

### Static and flow condition CFU analysis

We employed the procedural framework established by Joshi et al., (2025) (21). Silicone-coated latex catheter pieces (5 cm; commercial Foley catheter) were used as the biofilm substrate (supplementary Figure 2a, b). Bacterial cultures (0.1 OD₆₀₀) were introduced into catheter segments via 10 ml syringes and incubated statically for 4 hours at 37°C to permit initial adhesion. Catheter segments were then connected to sterile IV tubing lines attached to volumetric infusion pumps (flow rate: 200 µ l min⁻¹; Volume To Be Infused (VTBI): 288 ml over 24 hours). The inlet flask contained sterile MHB with or without sub-MIC antibiotic supplemented at the determined concentrations; the outlet flask was sealed with a sterile filter assembly to maintain sterility. Parallel static controls were prepared identically but without connected flow. After 24 hours, biofilm was recovered by flushing 2–3 mL of sterile 0.85% saline through the catheter segment and quantified by serial dilution plating on HiChrome UTI agar (37°C, 24 h), which permits chromogenic differentiation of *Enterococcus* spp. (teal colonies), *K. pneumoniae* (dark blue), and *P. aeruginosa* (green) on the same plate (supplementary Figure 3). CFU were calculated using CFU ml⁻¹ = (colonies counted × dilution factor)/volume plated (ml). Percentage survival for monocultures was calculated as the ratio of CFU under antibiotic treatment to CFU under the corresponding untreated control, within the same environmental condition: % Survival = (CFU_antibiotic ÷ CFU_untreated control) × 100. For polymicrobial communities, relative abundance of each species was calculated as its CFU divided by the summed raw CFU of all species present in that community, within the same condition: % Relative abundance = (CFU_species ÷ CFU_total) × 100. Additional biofilm was also collected from effluent passed through a sterile filter (filter fraction), permitting separation of catheter-associated (surface-attached) from dispersed (planktonic effluent) populations. Log₂ fold change was calculated relative to the corresponding static control condition for each species and combination to provide a normalized representation of pharmacological effect, and isolate flow- and antibiotic-specific effects from baseline growth. Heatmaps were plotted using GraphPad Prism 8.0.

### Microfluidic ibidi µ-slide VI ^0.4^ live/dead assay

For spatial and viability analysis, bacteria were seeded in channels of ibidi µ-slide VI ^0.4^ devices (ibidi, Gräfelfing, Germany) and allowed to adhere for 4 h at room temperature under static conditions (supplementary Figure 4). Channels were then connected to the volumetric infusion pump flow system at 200 µL min⁻¹ delivering sterile MHB with or without sub-MIC antibiotics for 24 h. After flow, channels were gently rinsed twice with sterile 0.85% saline. Live/dead staining was performed using the LIVE/DEAD BacLight kit (SYTO 9 and propidium iodide, PI; Thermo Fisher Scientific), prepared as 1:100 dilutions in sterile Milli-Q water, mixed 1:1, and applied (30 µL per channel) for 15 min at room temperature in the dark. Biofilms were imaged directly on the µ-slide using a confocal laser scanning microscope (Olympus laser scanning confocal microscope fluoview 3000) at 40×. SYTO 9 (live cells; intact membrane) was detected at excitation 498 nm, emission 517 nm; PI (dead/compromised cells) at excitation 590 nm, emission 610 nm. Z-stack images were acquired to characterize three-dimensional biofilm architecture. Cell counts were analysed using cell count plugin using ImageJ/Fiji software.

## Acknowledgement

The authors gratefully acknowledge the Department of Biotechnology (DBT), Government of India, for providing fellowship funding during the author (I.S) M.Sc. Biotechology teaching program phase II. We sincerely thank Ms. Pooja Gupta at the Vikram Sarabhai Institute of Cell and Molecular Biology (VSICMB), MSU for her expert technical assistance and support with confocal laser scanning microscopy (CLSM) imaging. We also thank the DBT -BUILDER funded microscopy facility at VSICMB. Author (R.M) acknowledges CSIR-UGC NET fellowship support from the Council of Scientific and Industrial Research and University Grants Commission, Government of India.

## Author contributions

**Ichha V Shah** (Conceptualization, Methodology, Investigation, Formal analysis, Data curation, Writing – original draft, Visualization), **Riddhin Modi** (Methodology, Investigation, Resources, Writing – review & editing) **Devarshi Gajjar** (Conceptualization, Resources, Supervision, Project administration, Funding acquisition, Writing – review & editing,)

## Data availability

Data will be made available upon request

## Ethics Approval

No ethical approval was required for this study.

## References

1. Cortese YJ, Wagner VE, Tierney M, Devine D, Fogarty A. 2018. Review of Catheter-Associated Urinary Tract Infections and In Vitro Urinary Tract Models. J Healthc Eng 2018:1–16.

2. Flores-Mireles AL, Walker JN, Caparon M, Hultgren SJ. 2015. Urinary tract infections: epidemiology, mechanisms of infection and treatment options. Nat Rev Microbiol 13:269–284.

3. Gade N, Burri R, Sujiv A, Mishra M, Pradeep BE, Debaje H, Sable T, Kaur A. 2023. Promoting Patient Safety: Exploring Device-Associated Healthcare Infections and Antimicrobial Susceptibility Pattern in a Multidisciplinary Intensive Care Units. Cureus 10.7759/cureus.50232.

4. Trautner BW, Darouiche RO. 2004. Role of biofilm in catheter-associated urinary tract infection☆. Am J Infect Control 32:177–183.

5. Jacobsen SM, Stickler DJ, Mobley HLT, Shirtliff ME. 2008. Complicated Catheter-Associated Urinary Tract Infections Due to Escherichia coli and Proteus mirabilis. Clin Microbiol Rev 21:26–59.

6. Gautam G, Satija S, Kaur R, Kumar A, Sharma D, Dhakad MS. 2024. Insight into the Burden of Antimicrobial Resistance among Bacterial Pathogens Isolated from Patients Admitted in ICUs of a Tertiary Care Hospital in India. Canadian Journal of Infectious Diseases and Medical Microbiology 2024:1–8.

7. Liu X, Kamperman M. 2025. Smart bacteria-responsive coatings for combating catheter-associated urinary tract infections. Mater Today Bio 34:102191.

8. Yuan L, Straub H, Shishaeva L, Ren Q. 2023. Microfluidics for Biofilm Studies. Annual Review of Analytical Chemistry 16:139–159.

9. Kim J, Park H-D, Chung S. 2012. Microfluidic Approaches to Bacterial Biofilm Formation. Molecules 17:9818–9834.

10. Yousefi Nojookambari N, Eslami G, Sadredinamin M, Vaezjalali M, Nikmanesh B, Dehbanipour R, Yazdansetad S, Ghalavand Z. 2024. Sub-minimum inhibitory concentrations (sub-MICs) of colistin on Acinetobacter baumannii biofilm formation potency, adherence, and invasion to epithelial host cells: an experimental study in an Iranian children’s referral hospital. Microbiol Spectr 12.

11. Gbejuade HO, Lovering AM, Webb JC. 2015. The role of microbial biofilms in prosthetic joint infections. Acta Orthop 86:147–158.

12. Choudhary P, Singh S, Agarwal V. 2020. Microbial BiofilmsBacterial Biofilms. IntechOpen. 10.5772/intechopen.90790

13. Kafil HS, Mobarez AM, Moghadam MF, Hashemi Z sadat, Yousefi M. 2016. Gentamicin induces efaA expression and biofilm formation in Enterococcus faecalis. Microb Pathog 92:30–35.

14. Samrot A V., Abubakar Mohamed A, Faradjeva E, Si Jie L, Hooi Sze C, Arif A, Chuan Sean T, Norbert Michael E, Yeok Mun C, Xiao Qi N, Ling Mok P, Kumar SS. 2021. Mechanisms and Impact of Biofilms and Targeting of Biofilms Using Bioactive Compounds—A Review. Medicina (B Aires) 57:839.

15. Stoodley P, Sauer K, Davies DG, Costerton JW. 2002. Biofilms as Complex Differentiated Communities. Annu Rev Microbiol 56:187–209.

16. Donlan RM, Costerton JW. 2002. Biofilms: Survival Mechanisms of Clinically Relevant Microorganisms. Clin Microbiol Rev 15:167–193.

17. Han A, Lee S-Y. 2023. An overview of various methods for in vitro biofilm formation: a review. Food Sci Biotechnol 32:1617–1629.

18. Pousti M, Zarabadi MP, Abbaszadeh Amirdehi M, Paquet-Mercier F, Greener J. 2019. Microfluidic bioanalytical flow cells for biofilm studies: a review. Analyst 144:68–86.

19. Hota S, Patil SR, Mane PM. 2024. Antimicrobial Resistance Profile of Enterococcal Isolates From Clinical Specimens at a Tertiary Care Hospital in Western Maharashtra, India. Cureus 10.7759/cureus.73416.

20. Wei Y, Palacios Araya D, Palmer KL. 2024. Enterococcus faecium: evolution, adaptation, pathogenesis and emerging therapeutics. Nat Rev Microbiol 22:705–721.

21. Joshi P, Bhattacharjee R, Sahu M, Gajjar D. 2025. Insights into urinary catheter colonisation and polymicrobial biofilms of Candida-bacteria under flow condition. Sci Rep 15:15375.

22. Sati H, Carrara E, Savoldi A, Hansen P, Garlasco J, Campagnaro E, Boccia S, Castillo-Polo JA, Magrini E, Garcia-Vello P, Wool E, Gigante V, Duffy E, Cassini A, Huttner B, Pardo PR, Naghavi M, Mirzayev F, Zignol M, Cameron A, Tacconelli E, Aboderin A, Al Ghoribi M, Al-Salman J, Amir A, Apisarnthanarak A, Blaser M, El-Sharif A, Essack S, Harbarth S, Huang X, Kapoor G, Knight G, Muhwa JC, Monnet DL, Ousassa T, Sacsaquispe R, Severin J, Sugai M, Taneja N, Umubyeyi Nyaruhirira A. 2025. The WHO Bacterial Priority Pathogens List 2024: a prioritisation study to guide research, development, and public health strategies against antimicrobial resistance. Lancet Infect Dis 25:1033–1043.

23. Elfadadny A, Ragab RF, AlHarbi M, Badshah F, Ibáñez-Arancibia E, Farag A, Hendawy AO, De los Ríos-Escalante PR, Aboubakr M, Zakai SA, Nageeb WM. 2024. Antimicrobial resistance of Pseudomonas aeruginosa: navigating clinical impacts, current resistance trends, and innovations in breaking therapies. Front Microbiol 15.

24. Sleziak J, Błażejewska M, Duszyńska W. 2025. Catheter-associated urinary tract infections in the intensive care unit during and after the COVID-19 pandemic. BMC Infect Dis 25:595.

25. Liu X, Sai F, Li L, Zhu C, Huang H. 2020. Clinical characteristics and risk factors of catheter-associated urinary tract infections caused by Klebsiella Pneumoniae. Ann Palliat Med 9:2668–2677.

26. Oleksy-Wawrzyniak M, Junka A, Brożyna M, Paweł M, Kwiek B, Nowak M, Mączyńska B, Bartoszewicz M. 2021. The In Vitro Ability of Klebsiella pneumoniae to Form Biofilm and the Potential of Various Compounds to Eradicate It from Urinary Catheters. Pathogens 11:42.

27. Kasetty S, Mould DL, Hogan DA, Nadell CD. 2021. Both Pseudomonas aeruginosa and Candida albicans Accumulate Greater Biomass in Dual-Species Biofilms under Flow. mSphere 6.

28. Li X, Lu N, Brady HR, Packman AI. 2016. Biomineralization strongly modulates the formation of Proteus mirabilis and Pseudomonas aeruginosa dual-species biofilms. FEMS Microbiol Ecol 92:fiw189.

29. Lehman SM, Donlan RM. 2015. Bacteriophage-Mediated Control of a Two-Species Biofilm Formed by Microorganisms Causing Catheter-Associated Urinary Tract Infections in an In Vitro Urinary Catheter Model. Antimicrob Agents Chemother 59:1127–1137.

30. Padron GC, Chen S, Sharma A, Modi Z, Koch MD, Sanfilippo JE. 2026. Shear flow promotes bacterial growth and shapes spatial gradients by rapidly replenishing scarce nutrients. mBio 17.

31. Rodesney CA, Roman B, Dhamani N, Cooley BJ, Katira P, Touhami A, Gordon VD. 2017. Mechanosensing of shear by Pseudomonas aeruginosa leads to increased levels of the cyclic-di-GMP signal initiating biofilm development. Proceedings of the National Academy of Sciences 114:5906–5911.

32. Nielsen H V., Guiton PS, Kline KA, Port GC, Pinkner JS, Neiers F, Normark S, Henriques-Normark B, Caparon MG, Hultgren SJ. 2012. The Metal Ion-Dependent Adhesion Site Motif of the Enterococcus faecalis EbpA Pilin Mediates Pilus Function in Catheter-Associated Urinary Tract Infection. mBio 3.

33. Guilhen C, Miquel S, Charbonnel N, Joseph L, Carrier G, Forestier C, Balestrino D. 2019. Colonization and immune modulation properties of Klebsiella pneumoniae biofilm-dispersed cells. NPJ Biofilms Microbiomes 5:25.

34. Zou Z, Pinkner JS, Obernuefemann CLP, Kleinschmidt KR, Sanick DA, Hickerson SM, Dodson KW, Henderson JP, Hultgren SJ, Caparon MG. 2026. Metabolic cross-talk promotes persistence of Enterococcus in a model of polymicrobial catheter-associated urinary tract infection. Sci Adv 12.

35. Ballén V, Ratia C, Cepas V, Soto SM. 2020. Enterococcus faecalis inhibits Klebsiella pneumoniae growth in polymicrobial biofilms in a glucose-enriched medium. Biofouling 36:846–861.

36. Todd K, Schneider O, Lawrence JM, Aronoff JL, Witek B, Velázquez-Colón V, Santana-Ufret V, Noda M, Relich RF, Zeng L, Limoli DH, Whidbey C, Vornhagen J. 2025. Pseudomonas aeruginosa phenazines dictate site-specific competitive interactions with Klebsiella pneumoniae 10.1101/2025.08.26.672413.

37. Holt JD, Schultz D, Nadell CD. 2024. Dispersal of a dominant competitor can drive multispecies coexistence in biofilms. Current Biology 34:4129–4142.e4.

38. Gibbs T, Levin SA, Levine JM. 2022. Coexistence in diverse communities with higher-order interactions. Proceedings of the National Academy of Sciences 119.

39. Laffont C, Wechsler T, Kümmerli R. 2024. Interactions between Pseudomonas aeruginosa and six opportunistic pathogens cover a broad spectrum from mutualism to antagonism. Environ Microbiol Rep 16.

40. Parthasarathy S, Jordan LD, Schwarting N, Woods MA, Abdullahi Z, Varahan S, Passos PMS, Miller B, Hancock LE. 2020. Involvement of Chromosomally Encoded Homologs of the RRNPP Protein Family in Enterococcus faecalis Biofilm Formation and Urinary Tract Infection Pathogenesis. J Bacteriol 202.

41. Zhang Y, Silva DM, Young P, Traini D, Li M, Ong HX, Cheng S. 2022. Understanding the effects of aerodynamic and hydrodynamic shear forces on Pseudomonas aeruginosa biofilm growth. Biotechnol Bioeng 119:1483–1497.

42. Cadavid E, Robledo SM, Quiñones W, Echeverri F. 2018. Induction of Biofilm Formation in Klebsiella pneumoniae ATCC 13884 by Several Drugs: The Possible Role of Quorum Sensing Modulation. Antibiotics 7:103.

43. Kumar A, Saha SK, Banerjee P, Prasad K, Sengupta TK. 2024. Antibiotic-Induced Biofilm Formations in Pseudomonas aeruginosa Strains KPW.1-S1 and HRW.1-S3 are Associated with Increased Production of eDNA and Exoproteins, Increased ROS Generation, and Increased Cell Surface Hydrophobicity. Curr Microbiol 81:11.

44. Bernardi S, Anderson A, Macchiarelli G, Hellwig E, Cieplik F, Vach K, Al-Ahmad A. 2021. Subinhibitory Antibiotic Concentrations Enhance Biofilm Formation of Clinical Enterococcus faecalis Isolates. Antibiotics 10:874.

45. Caixeta Magalhães Tibúrcio AA, Paiva AD, Pedrosa AL, Rodrigues WF, Bernardes da Silva R, Oliveira AG. 2022. Effect of sub-inhibitory concentrations of antibiotics on biofilm formation and expression of virulence genes in penicillin-resistant, ampicillin-susceptible Enterococcus faecalis. Heliyon 8:e11154.

46. Anderson CM, Mattenberger Y, Parga A, Portalier H, Tan CAZ, Esteban Henao M, Viollier PH, Kline KA. 2026. Enterococcus faecalis alters antibiotic susceptibility in Pseudomonas aeruginosa mixed-species biofilms. J Bacteriol 208.

47. Garratt I, Ravari MY, Clarke OE, McMurtrie J, Wand ME, Feil EJ, Taylor TB, Sutton JM, Jones B V. 2026. A model of polymicrobial catheter-associated urinary tract infection reveals biofilm-mediated modulation of treatment efficacy. J Appl Microbiol 137.

48. Founou RC, Founou LL, Allam M, Ismail A, Essack SY. 2024. Genome analysis of multidrug resistant Enterococcus faecium and Enterococcus faecalis circulating among hospitalized patients in uMgungundlovu District, KwaZulu-Natal, South Africa. BMC Infect Dis 24:671.

49. Galván EM, Mateyca C, Ielpi L. 2016. Role of interspecies interactions in dual-species biofilms developed in vitro by uropathogens isolated from polymicrobial urinary catheter-associated bacteriuria. Biofouling 32:1067–1077.

50. Webb JS, Thompson LS, James S, Charlton T, Tolker-Nielsen T, Koch B, Givskov M, Kjelleberg S. 2003. Cell Death in Pseudomonas aeruginosa Biofilm Development. J Bacteriol 185:4585–4592.

51. Saidi N, Davarzani F, Yousefpour Z, Owlia P. 2023. Effects of Sub-Minimum Inhibitory Concentrations of Gentamicin on Alginate Produced by Clinical Isolates of Pseudomonas aeruginosa. Adv Biomed Res 12.

52. Dale JL, Nilson JL, Barnes AMT, Dunny GM. 2017. Restructuring of Enterococcus faecalis biofilm architecture in response to antibiotic-induced stress. NPJ Biofilms Microbiomes 3:15.

53. Anderl JN, Franklin MJ, Stewart PS. 2000. Role of Antibiotic Penetration Limitation in Klebsiella pneumoniae Biofilm Resistance to Ampicillin and Ciprofloxacin. Antimicrob Agents Chemother 44:1818–1824.

54. Sinel C, Cacaci M, Meignen P, Guérin F, Davies BW, Sanguinetti M, Giard J-C, Cattoir V. 2017. Subinhibitory Concentrations of Ciprofloxacin Enhance Antimicrobial Resistance and Pathogenicity of Enterococcus faecium. Antimicrob Agents Chemother 61.

55. Rumbaugh KP, Sauer K. 2020. Biofilm dispersion. Nat Rev Microbiol 18:571–586.

56. Nye TM, Zou Z, Obernuefemann CLP, Pinkner JS, Lowry E, Kleinschmidt K, Bergeron K, Klim A, Dodson KW, Flores-Mireles AL, Walker JN, Wong DG, Desai A, Caparon MG, Hultgren SJ. 2024. Microbial co-occurrences on catheters from long-term catheterized patients. Nat Commun 15:61.

57. Dadashi Firouzjaei M, Hendizadeh P, Halaji M, Yaghoubi S, Teimourian M, Hosseini A, Rajabnia M, Pournajaf A. 2023. Quinolone Resistance in Biofilm-Forming Klebsiella pneumoniae-Related Catheter-Associated Urinary Tract Infections: A Neglected Problem. Jundishapur J Microbiol 16.

58. Gupta P, Chhibber S, Harjai K. 2016. Subinhibitory concentration of ciprofloxacin targets quorum sensing system of Pseudomonas aeruginosa causing inhibition of biofilm formation & reduction of virulence. Indian Journal of Medical Research 143:643–651.

59. Amer AN, Attia N, Baecker D, Mansour RE, El-Soudany I. 2025. Growth-Phase-Dependent Modulation of Quorum Sensing and Virulence Factors in Pseudomonas aeruginosa ATCC 27853 by Sub-MICs of Antibiotics. Antibiotics 14:731.

60. Mirzaii M, Alebouyeh M, Sohrabi MB, Eslami P, Fazli M, Ebrahimi M, HajiAsgarli P, Rashidan M. 2023. Antibiotic resistance assessment and multi-drug efflux pumps of Enterococcus faecium isolated from clinical specimens. The Journal of Infection in Developing Countries 17:649–655.

61. Shuppara AM, Padron GC, Sharma A, Modi Z, Koch MD, Sanfilippo JE. 2025. Shear flow patterns antimicrobial gradients across bacterial populations. Sci Adv 11.

62. Stewart PS, Camper AK, Handran, C.-T. Huang, M. Warnecke SD. 1997. Spatial Distribution and Coexistence of Klebsiella pneumoniae and Pseudomonas aeruginosa in Biofilms. Microb Ecol 33:2–10.

63. Cirz RT, O’Neill BM, Hammond JA, Head SR, Romesberg FE. 2006. Defining the Pseudomonas aeruginosa SOS Response and Its Role in the Global Response to the Antibiotic Ciprofloxacin. J Bacteriol 188:7101–7110.

64. Varga JJ, Zhao CY, Davis JD, Hao Y, Farrell JM, Gurney JR, Voit E, Brown SP. 2022. Antibiotics Drive Expansion of Rare Pathogens in a Chronic Infection Microbiome Model. mSphere 7.

65. Lebreton F, van Schaik W, Manson McGuire A, Godfrey P, Griggs A, Mazumdar V, Corander J, Cheng L, Saif S, Young S, Zeng Q, Wortman J, Birren B, Willems RJL, Earl AM, Gilmore MS. 2013. Emergence of Epidemic Multidrug-Resistant Enterococcus faecium from Animal and Commensal Strains. mBio 4.

66. Arias CA, Murray BE. 2012. The rise of the Enterococcus: beyond vancomycin resistance. Nat Rev Microbiol 10:266–278.

67. Gilmore MS, Salamzade R, Selleck E, Bryan N, Mello SS, Manson AL, Earl AM. 2020. Genes Contributing to the Unique Biology and Intrinsic Antibiotic Resistance of Enterococcus faecalis. mBio 11.

68. Willett JLE, Dunny GM. 2025. Insights into ecology, pathogenesis, and biofilm formation of Enterococcus faecalis from functional genomics. Microbiology and Molecular Biology Reviews 89.

69. Hallinen KM, Bodine SP, Stone HA, Muir TW, Wingreen NS, Gitai Z. 2025. Bacterial species with different nanocolony morphologies have distinct flow-dependent colonization behaviors. Proceedings of the National Academy of Sciences 122.

70. Antypas H, Schmidtchen V, Staiger WI, Yanhong L, Tan RJW, NG KKF, Neo CJY, Radhesh SM, Tanoto FR, da Silva RAG, Colomer-Winter C, Schütz SD, Kloehn J, Muthualagu Natarajan L, Manzano C, Wong JJ, Pethe K, Hasse B, Brugger SD, Wong SL, Van Tyne D, Zinkernagel AS, Kline KA. 2026. Loss of Fsr quorum sensing promotes biofilm formation and worsens outcomes in enterococcal infective endocarditis. Nat Commun 17:1668.

71. Gaston JR, Andersen MJ, Johnson AO, Bair KL, Sullivan CM, Guterman LB, White AN, Brauer AL, Learman BS, Flores-Mireles AL, Armbruster CE. 2020. Enterococcus faecalis Polymicrobial Interactions Facilitate Biofilm Formation, Antibiotic Recalcitrance, and Persistent Colonization of the Catheterized Urinary Tract. Pathogens 9:835.

72. Stepanović S, Vuković D, Dakić I, Savić B, Švabić-Vlahović M. 2000. A modified microtiter-plate test for quantification of staphylococcal biofilm formation. J Microbiol Methods 40:175–179.

